# GW: ultra-fast chromosome-scale visualisation of genomics data

**DOI:** 10.1101/2024.07.26.605272

**Authors:** Kez Cleal, Alexander Kearsey, Duncan M. Baird

## Abstract

Genome-Wide (GW) is an interactive genome browser that expedites analysis of aligned sequencing reads and data tracks, and introduces novel interfaces for exploring, annotating and quantifying data. GW’s high-performance design enables rapid rendering of data at speeds approaching the file reading rate, in addition to removing the memory constraints of visualizing large regions. We report substantial gains in performance and demonstrate GW’s utility in exploring massive genomic regions or chromosomes without requiring additional processing.

---

Genome browsers such as the Integrative Genomics Viewer (IGV) and JBrowse2 (1, 2) are invaluable for exploring aligned sequencing reads and verifying variant calls, but lack the speed offered by high-performance libraries and frameworks. In particular, current tools struggle to visualise large genomic regions which is a common need when investigating complex chromosomal rearrangements that can encompass megabases (3, 4). Exploring collections of variants and manually annotating data remains cumbersome due to slow loading times or requiring static image generation.

Here we describe GW, an interactive browser for rapidly visualising aligned sequencing reads, data tracks and genome-variation datasets, featuring a multithreaded architecture for desktop or command-line use on multiple platforms, including cloud environments (e.g. AWS servers or the Genomics England Research Environment) (Fig. 1a-c, Supplementary Movie1.mp4, Supplementary Table1).

**Fig 1.**
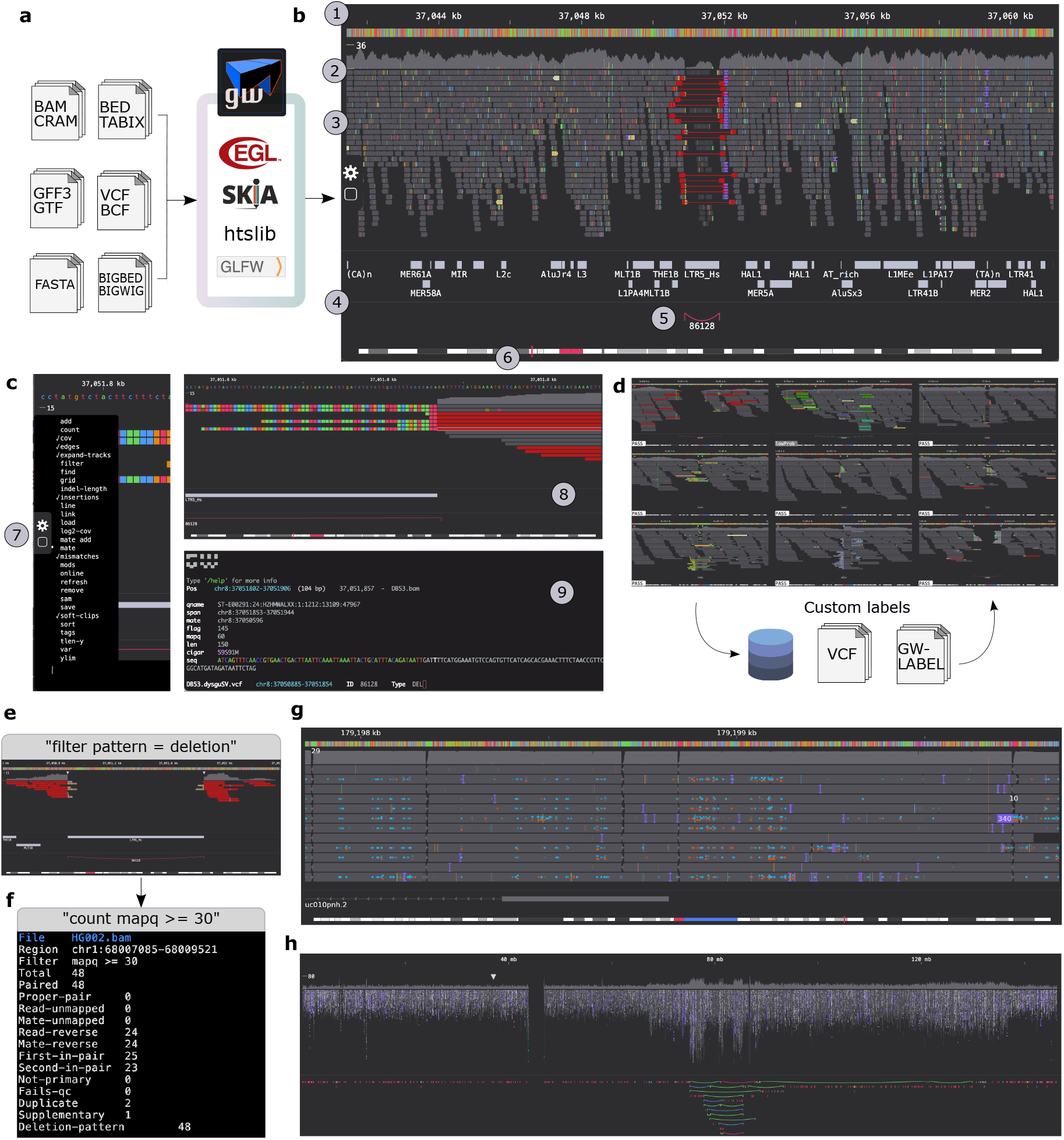
Genome browsing and variant annotation using GW. **(a)** GW accepts common data formats and uses libraries such as Skia and htslib for faster visualization. **(b)** The alignment-view displays a reference genome with scale bar (1), coverage and SNP profiles (2), stacked reads (3), and data tracks below (4). An SV is indicated with a colored arc (5). An ideogram is overlayed with a box to indicate chromosome location that can be used as a slider to change position (6). **(c)** Access commands and functions via the bubble (6) or the command box (7). Data interactions (8) display information to the terminal (9). **(d)** The image-view displays a grid of images, enabling manual annotation. GW supports versatile filtering **(e)** and counting **(f)** commands and supports display of multiple base modifications simultaneously **(g). (h)** Dynamic visualization of chr8 (146Mb), showing a complex rearrangement pattern and copy-number profile, where the white arrow shows the location of **(a)**.

GW supports extensive data interactions using 35 built-in commands, enabling users to load, save, navigate and search files, filter and count reads, change appearances and organise data (Fig. 1e-g). Data is displayed using either the ‘alignment-view’ or ‘image-view’. With the alignment-view (Fig. 1b), sequencing reads are arranged as stacks with colours representing information such as mismatches or discordant reads. Coverage profiles appear above and users can arrange multiple files and regions in rows and columns. Data tracks are placed below, and can be expanded, collapsed, and dynamically resized. Interactive elements display information to the terminal (Fig. 1c). GW supports themes including ‘slate’ (shown) and ‘dark’ to reduce eye strain, and ‘igv’ (Supplementary Fig. 1) (1).

The ‘image-view’ (Fig. 1d) organises images into a grid, simplifying exploration and curation of large datasets. Images can be loaded from disk or generated in real-time from files, while right-clicking an image will switch between alignment and image-views, permitting static-image exploration and dynamic data interactions. Multiple input files are supported for comparing variant sets and manual curation is facilitated by customizable labels or information parsed from input VCF’s. Labels are displayed over thumbnail images, and can be cycled, tagged with comments, loaded from disk, or encoded within files (Fig. 1d).

GW can create static images from variant files, reference genome indexes, or genome coordinates, and can save images of every chromosome in <1min30s (Supplementary Fig. 2). Images can later be explored as filenames encode genome locations. Loading of chromosome-images alongside bam files (Supplementary Fig. 3), permits right-clicking on a chromosome-image which switches the user to the alignment-view, zooming into a 500kb genome region for rapid data exploration.

GW also introduces new tools for data analysis not available in other browsers, including composable filtering and counting commands, allowing for tasks such as counting the number of reads that support a structural variant (SV), eliminating the need for manual counting or extra tools (Fig. 1e, f). Eight additional analysis tasks are provided as examples (Supplementary Fig. 4), including filtering/counting reads containing kmers (e.g. ‘count seq contains TTAGGG’ or ‘filter seq omit AAAAAA’), regional-overlaps (‘filter pos >= 10000 and ref-end < 20000’), or sam-tag properties (‘count AS > 100’, ‘filter NM = 0’), which would be difficult or impossible using other browsers. Additional display modalities in GW include drawing modified bases, representing SV patterns using coloured arcs (Fig. 1g, h), and scaling read-position by template-length and (Supplementary Fig. 5).

GW links to online databases using the ‘online’ command, providing web-links to UCSC, Gnomad, Ensemble, NCBI and DECIPHER browsers with the matching genome location as seen in GW, making it straightforward to compare samples against data tracks and population or disease databases.

We compared GW’s speed to other browsers like IGV and JBrowse2 (jb2export plugin), along with static image tools like samplot (1, 2, 5–7). GW generated images close to the file reading speed (Supplementary Fig. 6), significantly outpacing other tools on all datasets (Supplementary Fig. 7-12, Supplementary_Data.xlsx). For example, compared to IGV using a 2 Mb region size, GW showed >146-fold improvement in rendering speed (median 185-fold), while also using up to 109-fold less memory (Illumina data). Using a smartphone, GW was >30× quicker than IGV on a desktop. For large-scale analysis, GW visualized chr8 (146Mb, 26× coverage) from a sample with a complex rearrangement pattern (4) in 7.2 seconds using 736Mb of memory, showcasing its efficiency in processing entire chromosomes without additional steps (Fig. 1h).

Future work will focus on adapting GW for rendering alignments against pangenome graphs. In summary, GW permits rapid exploring, analysing and annotating genomic data on a much larger scale.

## Online Methods

### Fast data rendering

GW relies on several high-performance libraries to achieve fast data rendering, including htslib (8) for alignment file reading, Skia for fast 2D vector graphics rendering (9), and GLFW for window management (10). GW is written in C++ and has an efficient design, where for smaller regions minimal data is buffered in memory, while for larger regions, alignment data can be streamed without buffering.

GW employs various Level Of Detail (LOD) thresholds according to the region size being displayed, above which certain image details are not drawn, or elided. These user-customisable threshold are designed to both increase the clarity of images and speed up rendering. For example, above a region size of 20 kb, the reference track is not drawn, whilst above 500 kb, mismatches are not drawn. Another method to speed up drawing is using gap elision, where deletion-gaps in alignments are not drawn if they are too small to be visualised on-screen, occurring when their size falls below a threshold (1 + region-size / image-width), where width is measured in pixels and region-size is in base-pairs.

### Window management and graphics

GW utilizes the GLFW library for managing user inputs via the keyboard, mouse or other devices like a touchpad, and managing the interface between the operating system window and a graphics surface which can reside in GPU memory or main memory. For drawing, GW uses the Skia graphics library from Google, which translates drawing calls to a target graphics API, and manages drawing batches. For systems other than MacOS, Skia is built with the EGL backend allowing the use of the OpenGL ES API, whilst for MacOS the native OpenGL API is utilised.

When rendering a static image, GW creates a graphics context in main memory rather than on the GPU which allows for efficient parallel drawing between multiple threads and multiple contexts. When rendering to screen, surfaces are created on both the GPU and CPU side. Most rendering is carried out on the CPU side using a raster surface. As most draw operations consist of simple rectangles, these are efficiently handled by the CPU, and we found transporting these commands to the GPU incurred a non-trivial time penalty which was large enough to warrant performing rendering using a raster surface. Once the frame has been rendered to the raster surface, the image is transferred to the GPU surface for display using the GLFW interface.

Each frame rendered by GW is cached, allowing GW to redraw only sections of the screen that have experienced on update since the last frame. For example, for multi-sample or multi-region views, updates to one screen panel do not invoke re-processing or redrawing of other panels which speeds up the frame rate.

### Finding Y-position of reads

The on-screen y position, or row, of reads is determined before drawing and can take an integer value in the range [-1, ylim], where ylim defaults to 60 but can also be set by the user. Reads that are out-of-view will have ylim = -1. To determine the on-screen position of reads, an end-array is used with length equal to ylim, with each value representing the reference end position of each row. For each incoming read, the y-position is set as the index in the end-array whose value is less than the alignment position. The value in the end-array is then updated to reflect the reference-end position of the new read. This operation is used to append reads to the right-hand-side of the read queue. To append reads to the left-hand-side of the read queue a separate start-array is used (length = ylim) that performs the opposite function, keeping track of the start position of each row of reads. Left-appending reads would ideally require input reads to be sorted by reference-end position, although reading the alignment file returns reads sorted by reference start. To avoid having to sort input reads, reads are appended to the left-hand side in reverse order. Using these arrays allows for appending new reads to either side of the read queue in linear time without needing to re-analyse buffered data.

### Fast scrolling

To speed up drawing GW buffers in-view data only, up until a threshold region size is reached which is 1.5Mb by default. Above this threshold then the entire frame is rendered by streaming data from disk which removes the memory constraints of visualising large regions. However, below this threshold, to speed up scrolling, GW reuses buffered reads from the previous frame and only loads reads from newly requested regions, whilst avoiding fetching reads from out-of-view loci, which reduces file loading pressure. Newly fetched data is added to the left or right-hand-side of the read queue, maintaining position sorted-order and any out-of-view data is dropped. Zooming-out involves appending reads to both sides of the queue, whilst zooming-in involves culling reads from both sides of the queue. To ensure stacks of alignments remain in the same position on-screen during scrolling and zooming, the y-positions of new read alignments are determined using the end-array and start-array.

### Grid layout and image generation

GW can display images in a grid layout by loading files from the filesystem (png format), or generating images on the fly from a VCF/BCF or BED files. Files loaded from disk are processed in parallel. Images are stored in memory making scrolling to previously-seen images much faster. If the window size changes or there is a change to image settings, or the refresh command is used, buffered images are discarded.

### Filtering and counting reads

GW supports ‘filter’ and ‘count’ commands that can accept a simple expression for filtering and quantifying aligned sequencing reads. Reads can be filtered or counted using an expression of the form ‘{property} {operation} {value}’, where a ‘property’ relates to the alignment, ‘operation’ is a standard operator, and ‘value’ is the desired filtering value. Currently, over 40 different properties can be used, derived from attributes of the alignment (mapq, pos, tlen etc), flag values (‘paired’, ‘proper-pair’ etc or numeric values, 1, 2, respectively), or standard SAM-tag values (Alignment score AS, Edit distance NM etc). These properties can be combined with up to 10 operators which include common comparison operators (greater than ‘>‘, equal to ‘==‘ etc) as well as bitwise ‘&’ operator, and sequence specific operators (‘contains’ and its opposite ‘omit’). The value side of the expression is chosen by the user and is typically a numeric value, but can also be a string value. Expressions can also be chained to create more complex subsets of the data and may be useful for many analytical tasks that would normally require additional bioinformatics tools. To illustrate some brief examples, sub setting the data by mapping quality, insert size or location-of-mate is possible, and more complex analysis such as counting the intersection of reads with a genome feature, or filtering/selecting reads by the occurrence of a kmer is also possible.

### Benchmarking

We benchmarked visualisation tools using paired-end Illumina data from the HG002 human sample sequenced to 40X coverage (148bp reads), and PacBio HiFi data at around 8X. Reads were aligned to the ucsc.hg19 reference using bwa mem (v0.7.17, paired-end reads), or minimap2 (v2.21 long-reads) (11). The benchmark scripts (benchmark.py and plot.py) and results are available online in the GW repository at https://github.com/kcleal/gw/tree/master/test. GW (v1.0.3), IGV (v2.16.0), JBrowse2 (v2.4.0, jb2export “img” plugin), samplot (v1.3.0), and genomeview (v1.1.4) were invoked from the command line to generate a single image of a target region. Regions that overlapped gaps in the reference genome were ignored, as were regions with zero coverage. Sampled regions (n=20) were the same for each application. All tools were run in single-thread mode, while GW and IGV were also tested using 4 threads. IGV was configured using the desktop interface to display large regions by setting the “Visibility range threshold to 250Mb”. Other parameters were configured to try and make a fair comparison with GW, turning off “Quick consensus mode”, “Compute insert size thresholds”, “Show insertion markers”, “Downsample reads”, whilst it was ensured the following options were turned on “Show mismatched bases”, “Show soft-clipped bases”. Mean elapsed wall time was measured using hyperfine (12), and memory (resident set size) were measured using the GNU-time application. The time taken to visualize a 2bp region was taken as the initialisation time of the application, which included task such as file opening, saving of the image and application shutdown. For larger regions, the additional time taken on top of this was taken as the render time of the application. The maximum speed that the alignment file could be read was approximated as the average time to run “samtools view -c -@{t}” to count the number of reads in the region, employing the same number of threads as the comparison visualization tool, setting {t} to either 1 or 4. The benchmark was run on an Intel i9-11900K (Iris graphics), NVMe WD 2TB, 64 GB memory. If a tool took more than 3 hours to complete the benchmark, it was not included in the results. For displaying modified bases, GW was compared to IGV as other tools either did not support modified bases, or we were unable to configure the tool to do so using their command-line interface.

Benchmarking of GW was also performed on a Google Pixel 6 smartphone, using HG002 40X Illumina sample. Here, the benchmark was limited to chromosome 1 due to space limitations and execution time was measured using hyperfine (12). Movie1.mp4 was generated on a MacBook Pro M2 16 Gb memory.

## Data Availability

The HG002 human sample (13), sequenced using Illumina (40×coverage), was downloaded from GIAB ftp://ftp-trace.ncbi.nlm.nih.gov/giab/ftp/data/AshkenazimTrio/HG002_NA24385_son/NIST_HiSeq_HG002_Homogeneity-10953946/HG002Run01-11419412/HG002run1_S1.bam). PacBio HiFi data for HG002 (8× coverage) was downloaded from SRA (https://www.ncbi.nlm.nih.gov/sra) under accession SRR10188368. ONT data from HG002, sequenced using the R9.4.1 HAC protocol, was downloaded from SRA accession SRR11537600. Additionally, ONT data from HG002, sequenced using the “Kit14” workflow with methylation base calls, was downloaded from AWS s3://ont-open-data/giab_2023.05/analysis/hg002/sup/, sample PAO89685. A synthetic high-coverage sample was created by duplicating the PAO89685 sample. The benchmark script and results can be found online in the GW repository at https://github.com/kcleal/gw/tree/master/test. Sample DB53 has been reported previously in (4), and is available from NCBI BioProject PRJNA417592.

## Availability of supporting source code and requirements

Project name: GW (Genome Wide)

Project homepage: https://github.com/kcleal/gw

Operating system(s): unix based, Windows.

Programming language: C++

Other requirements: glfw, htslib

License: MIT

## Abbreviations

IGV: Integrative Genome Viewer
GIAB: Genome In A Bottle consortium
GPU: graphics processing unit
NVMe: non-volatile memory express
SV: structural variant
ONT: Oxford Nanopore Technologies
AWS: Amazon Web Services

## Conflict of interest disclosure

The authors declare that they have no competing interests.

## Funding

Work in the Baird laboratory is funded by Cancer Research UK (C17199/A29202) and the Wales Cancer Research Centre.

## Author contributions

KC designed and wrote the software, performed experiments, and wrote the manuscript. AK contributed code, help create distributions and design ideas, and tested the software. DMB provided feedback and performed manuscript editing. All authors read and approved the final manuscript.

## Notes

### Competing Interest Statement

The authors have declared no competing interest.

### Summary of Updates

The version updates the benchmarking results to GW v1.0.3. Importantly, this version corrects an important bug that affected the benchmarking of smaller regions (displayed in Supplementary Figures 6-12), causing an over estimation of GW's performance. The largest region size was unaffected by the bug.

