## Supplementary Fig for "GW: ultra-fast chromosome-scale visualisation of genomics data"

**Supplementary Movie 1. Exploring the genome using the alignment view.** Demonstration of the GW desktop app, showcasing the scrolling and zooming speed of GW, and exploration of large copy number variants and structural-variant calls.

| Features | IGV | JBrowse2 | GW |
| --- | --- | --- | --- |
| 1 Alignment view | *** | *** | *** |
| 2 Image view | - | - | *** |
| 3 Command-line usage | * | ** | *** |
| 4 Fast start-up time | * |  | *** |
| 5 Rendering performance | * | * | *** |
| 6 Simple interface | ** | * | *** |
| 7 Raw data accessibility | * | * | *** |
| 8 Fast zooming and scrolling | * | * | *** |
| 9 Data types supported | *** | *** | ** |
| 10 Manual annotation | - | - | *** |
| 11 Filtering and quantification of alignment data | * | * | *** |
| 12 Variant exploration | ** | *** | *** |
| 13 Remote access | ** | ** | *** |

**Supplementary Table 1. Comparison of features between genome browsers.**

1. All browsers implement first class support for viewing alignment data in BAM/CRAM format, showing stacks of reads against a linear reference genome along with coverage profiles, with colours representing additional information such as mismatches or discordant reads.
2. GW uniquely supports viewing static images using a grid layout. These can be generated on the fly or loaded from disk. By encoding genome coordinates in filenames, this feature enables users to re-open a static image for dynamic viewing, providing a convenient and fast way to revisit data views encoded as images. One interesting use of this feature is being able to generate chromosome-scale images which can be dynamically viewed in a fast manner.
3. GW can be used from the command-line whilst other tools have more limited support. IGV can be used via a batch script and JBrowse2 has a number of plugins that can be utilised, although neither offers the convenience of GW.
4. IGV and JBrowse2 take a significant amount of time to start the application and load data, particularly JBrowse2 which requires manual loading of reference genome, alignment data and other data tracks. GW starts in < 0.2s and users can drag-and-drop data files into the window. IGV also supports drag-and-dropping.
5. IGV and JBrowse2 can render small regions of data quickly, but become slow when viewing large regions. GW can render much large regions, up to 20 Mb per second.
6. GW has a very simple interface that is intuitive to use. Whilst IGV and JBrowse21 have deep customisation options, their interface requires more time to learn.
7. GW can display raw (uncompressed) data to the terminal (FASTA, BAM, VCF, BED) which is not always possible using other tools which tend to display representations of the data. Raw data can be useful for some bioinformatics tasks, for example, printing the reference genome sequence at a specific locus is achieved in GW by simply clicking on the reference track, resulting the sequence being printed to the terminal.
8. Zooming and scrolling are much faster in GW using mouse or keyboard keys making data tracks much quicker to explore on a wider scale.
9. IGV and JBrowse2 supports a large number of datatypes. GW supports fewer, but still covers many common data formats including (BAM/CRAM, FASTA, BED, BIGBED, BIGWIG, VCF/BCF, TABIX, GFF3, GTF).
10. GW has first-class support for manually annotating variants, using customizable labels, or labels parsed from input files. Annotations can be updated over multiple sessions.
11. IGV and JBrowse2 have basic filtering options, for example, reads can be filtered by mapping quality of alignment tag. GW supports a more powerful filtering interface that can be used to select reads with particular attributes, locations, or sequence content. GW can also efficiently count and categorize reads for data analysis which is not possible using other tools.
12. IGV can load data into a region explorer or cycle through variant in a VCF/BCF file using hotkeys. JBrowse2 implements a feature rich table browser for variant data. GW displays variants using thumbnail images in a grid which can help provide high-level overviews of data types such as structural variants, and convey additional patterns in the data.
13. IGV and JBrowse2 can both be used over remote connections, and JBrowse2 has the ability to be embedded within webpages. GW can be quickly launched from the command-line with X11-forwarding enabled, normally requiring only a single command-line argument (the BAM file), making GW the most convenient to use on remote servers.

**a**

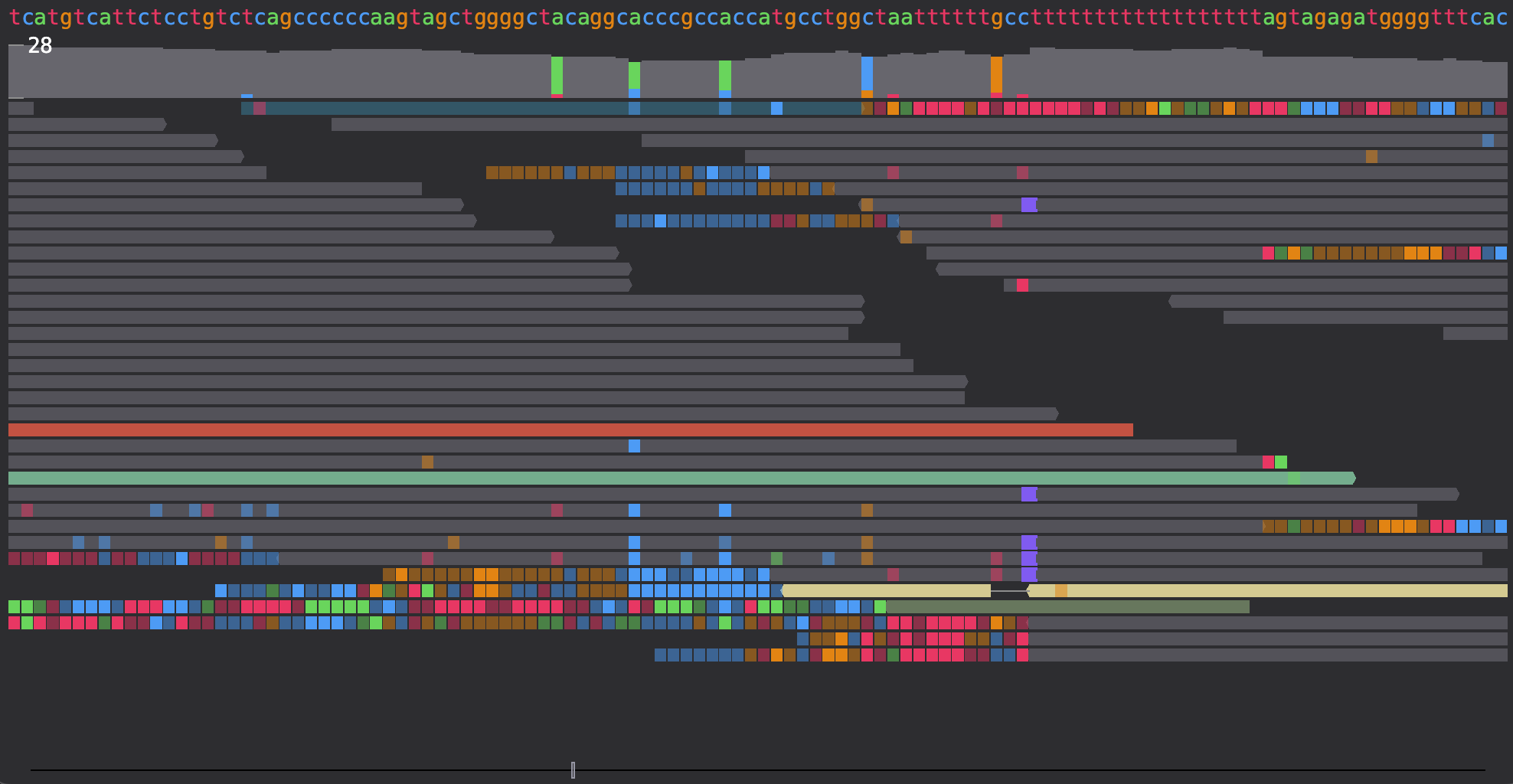

**b**

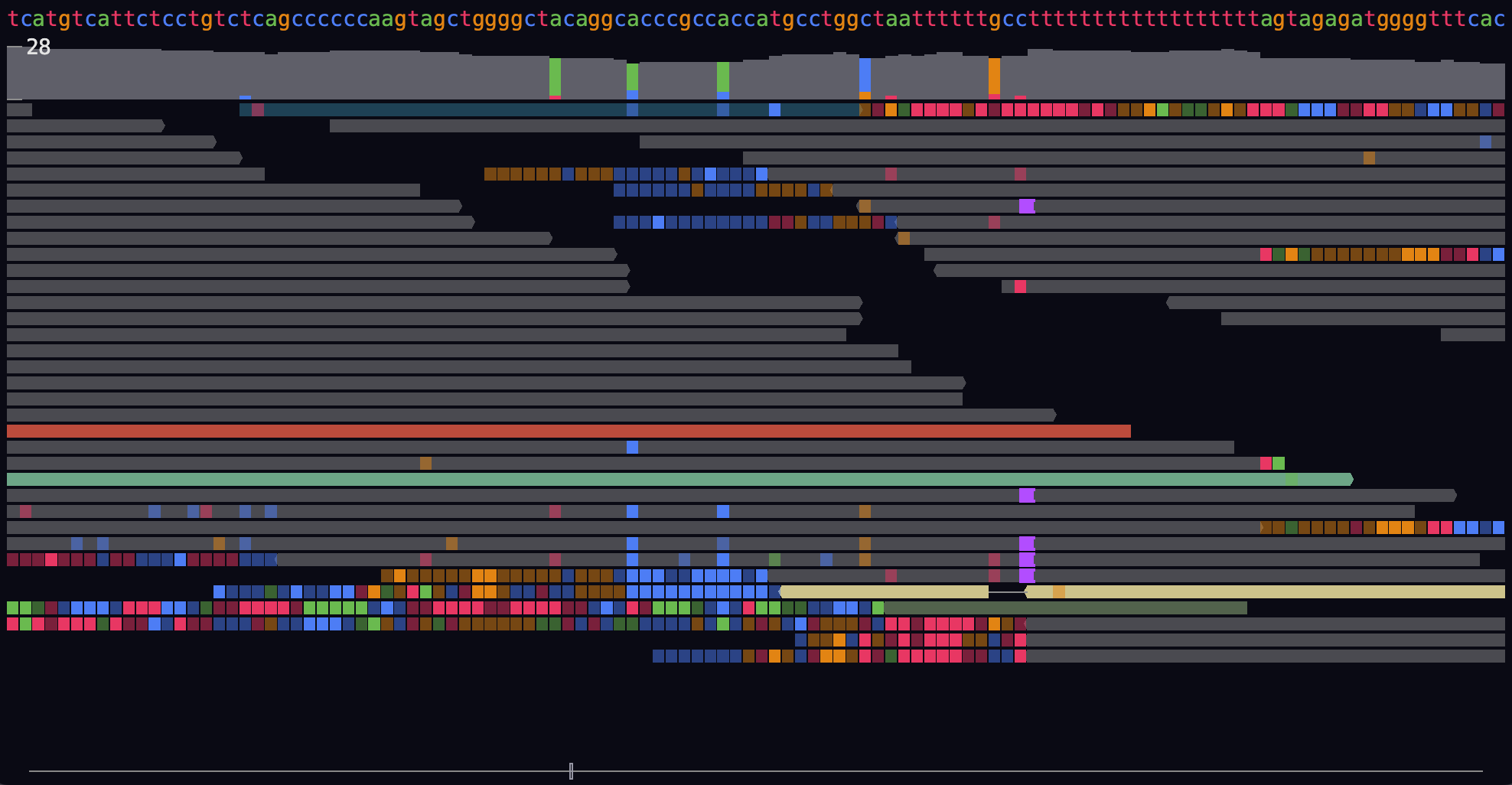

**c**

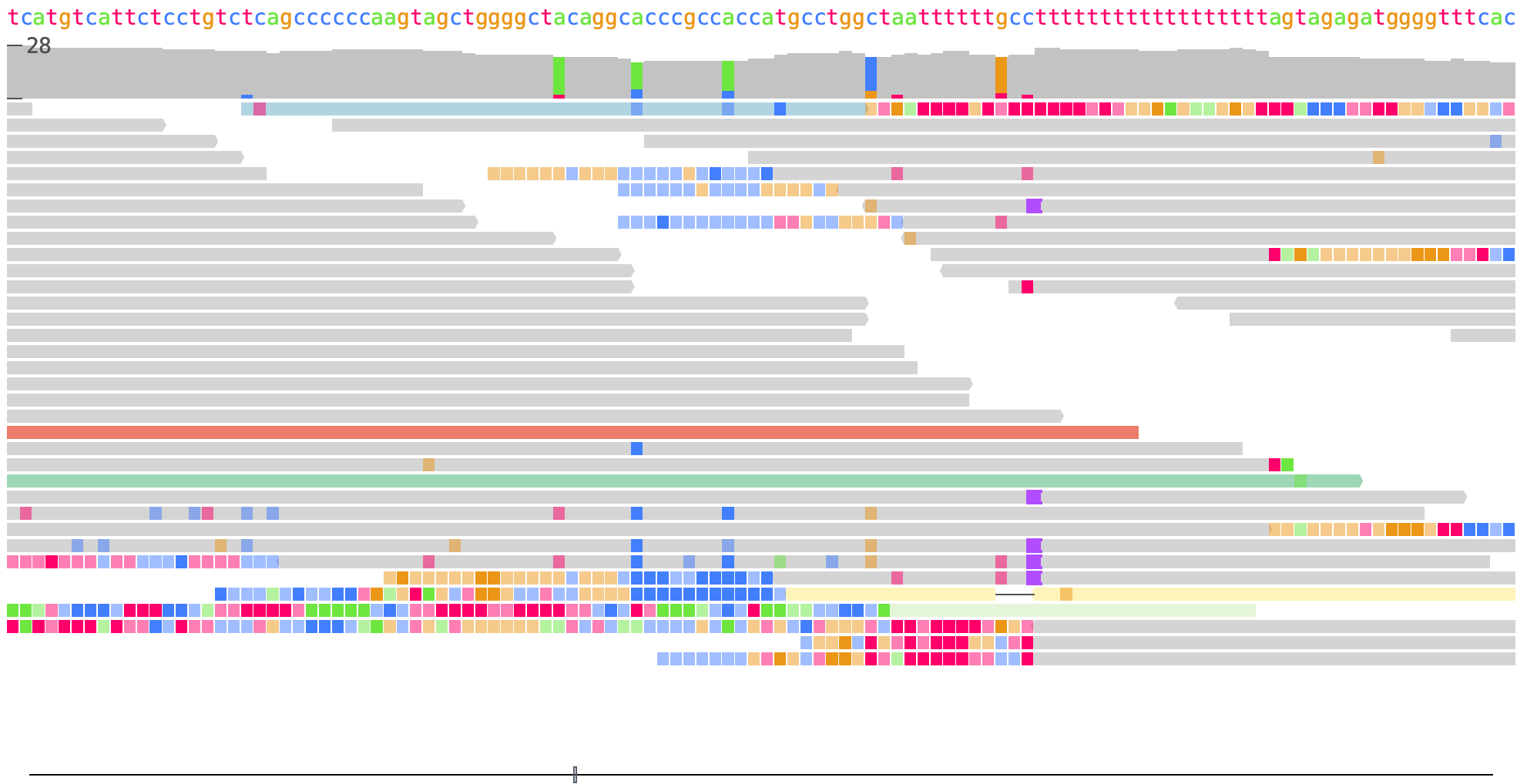

**Supplementary Figure 1. GW themes.** The slate (a) and dark (b) themes are designed to reduce eye-strain, whilst the igv (c) theme may also be used.

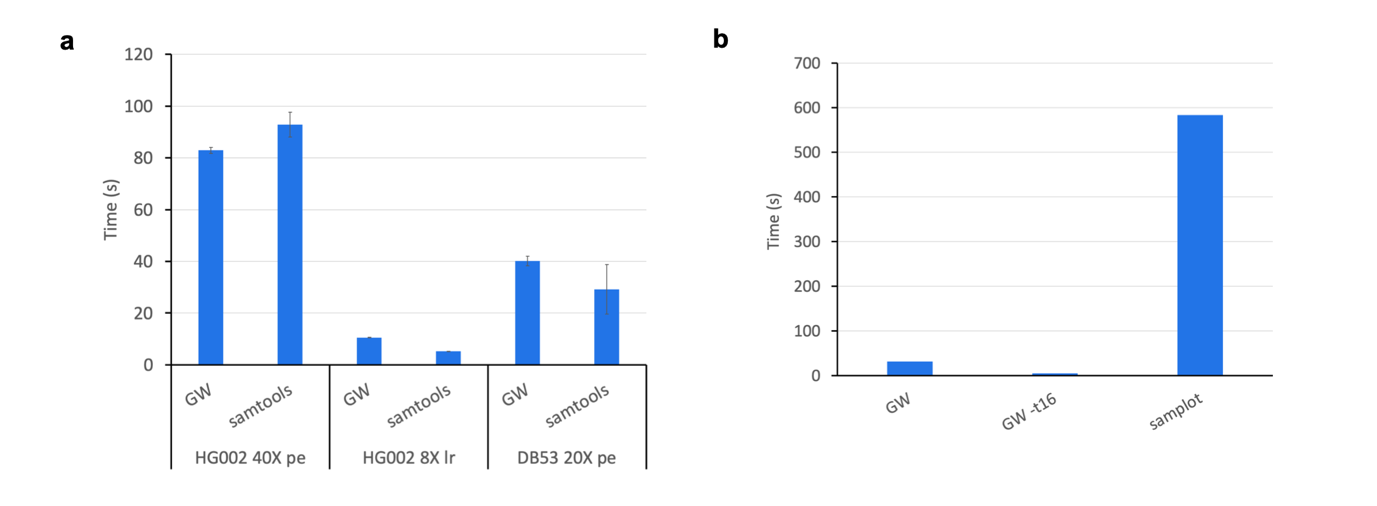

**Supplementary Figure 2. Image generation from genome index or VCF file.** The time taken to render chromosome images from a whole BAM file using 16 threads was assessed in (a) and compared to the time taken to use samtools to count reads reads, also using 16 threads. The time taken to generate 1000 images from a VCF was assessed (b), comparing against samplot only (-t16 indicates 16 threads were used, b). pe – paired-end, lr – long reads, X refers to the coverage of the sample.

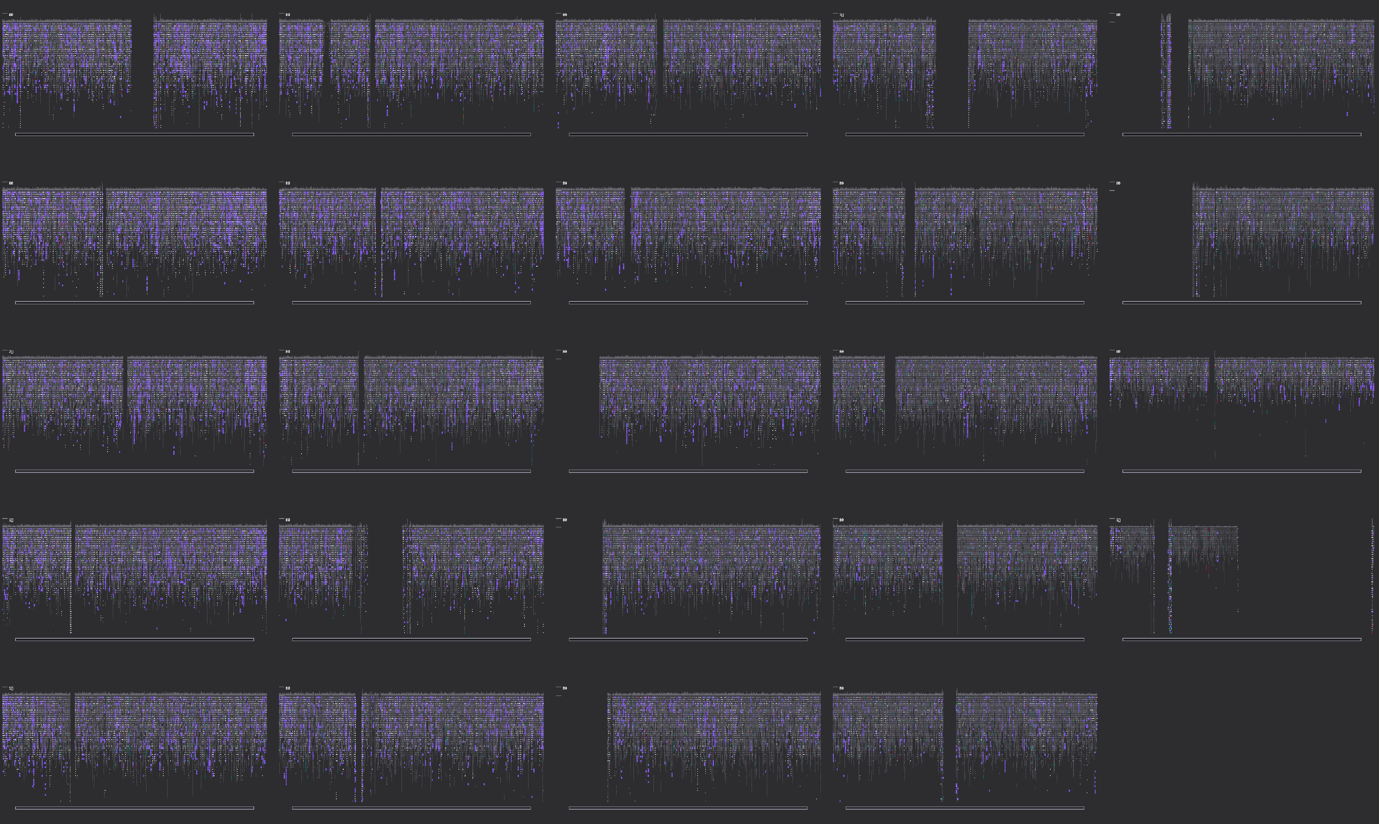
**Supplementary Figure 3. Chromosome plots opened using the image view.** A. All 24 canonical chromosomes are displayed using the image view. Chromosome labels have been added manually.

chrY

chrX

chr22

chr21

chr20

chr19

chr18

chr17

chr16

chr15

chr14

chr13

chr12

chr11

chr10

chr9

chr8

chr7

chr6

chr5

chr4

chr3

chr2

chr1

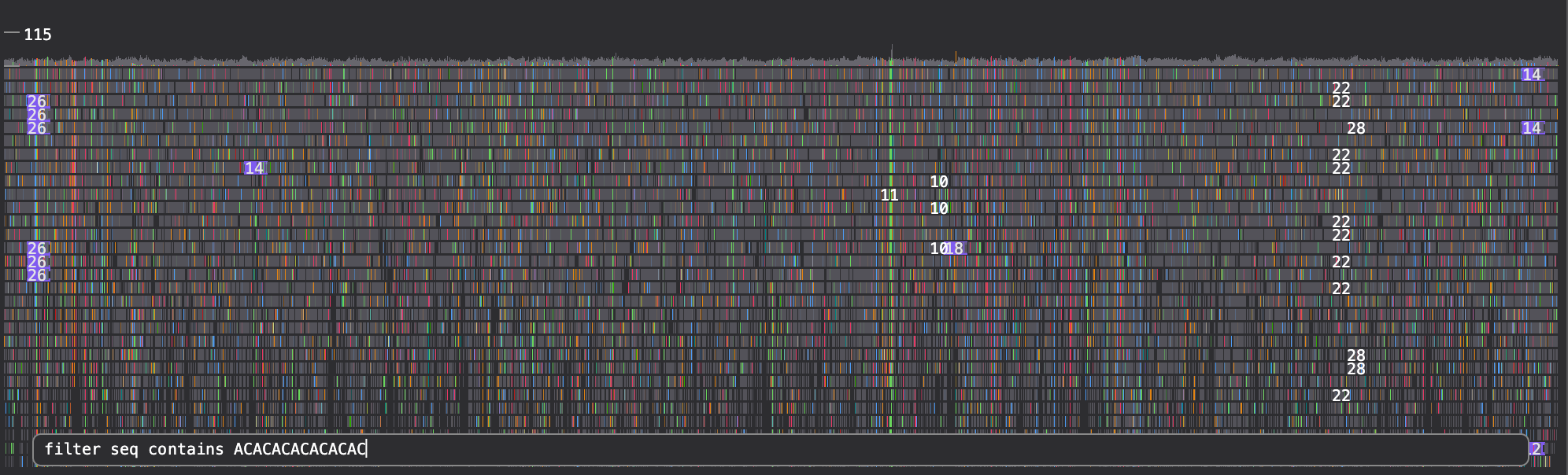

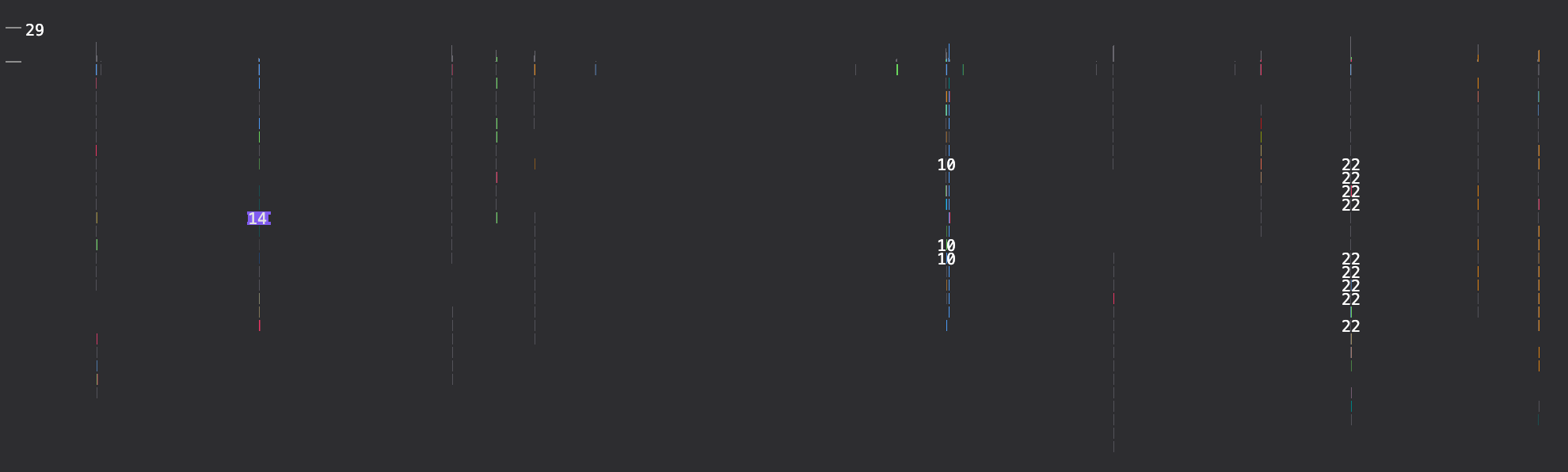

Identify reads containing kmer of interest

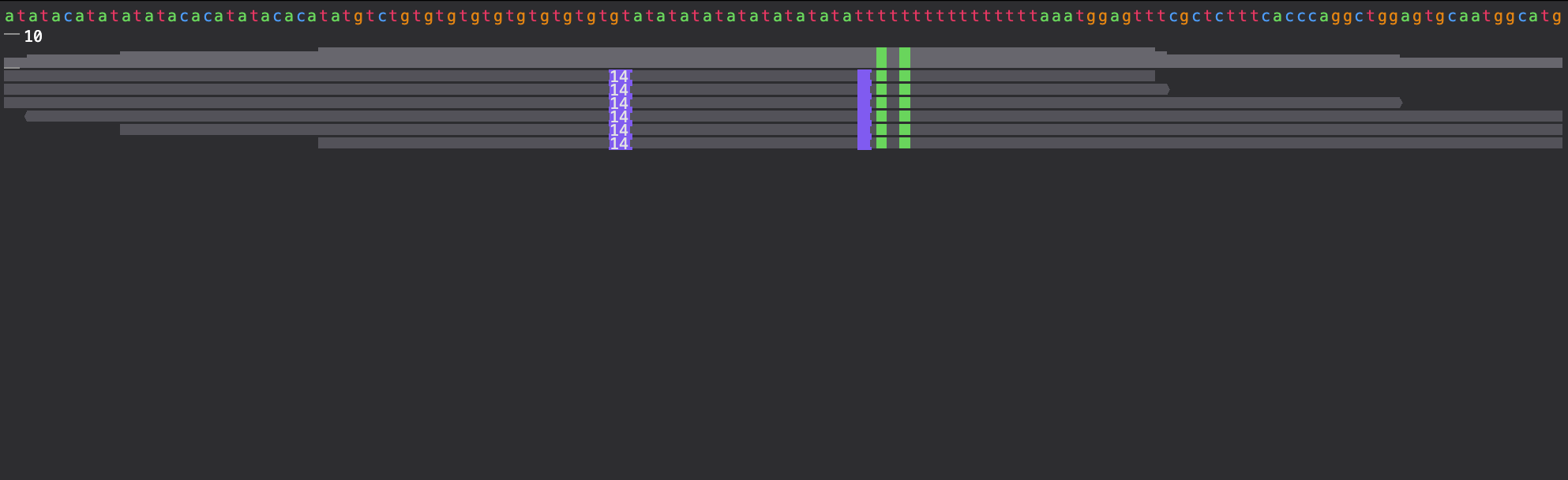

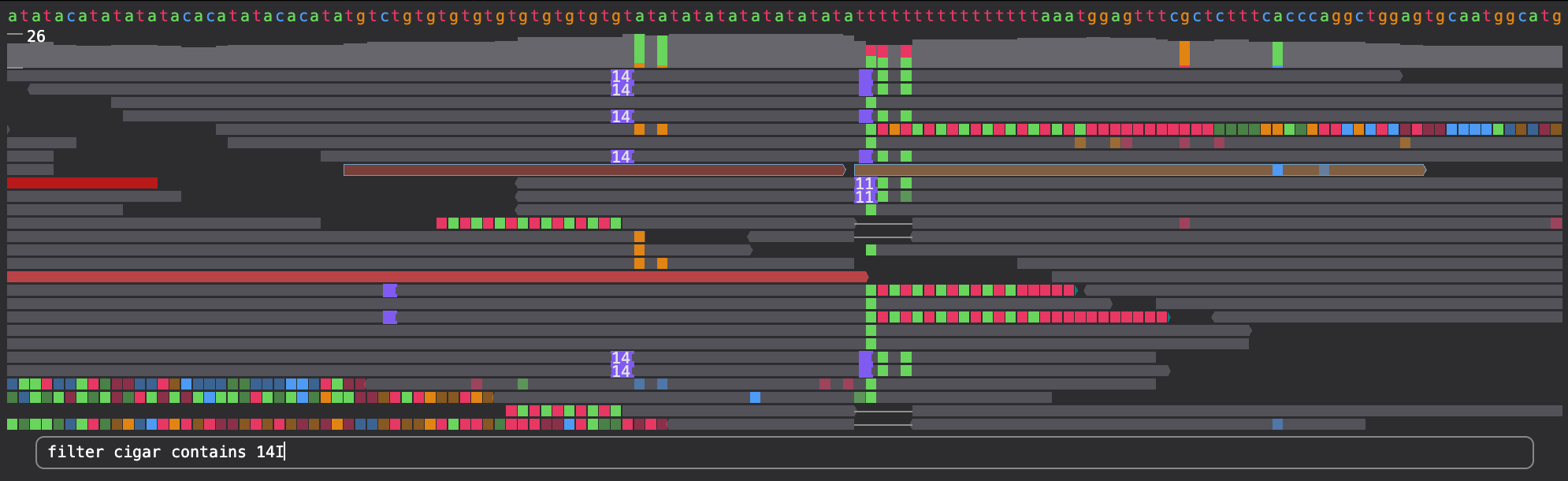

Identify reads with a particular cigar pattern “14I”

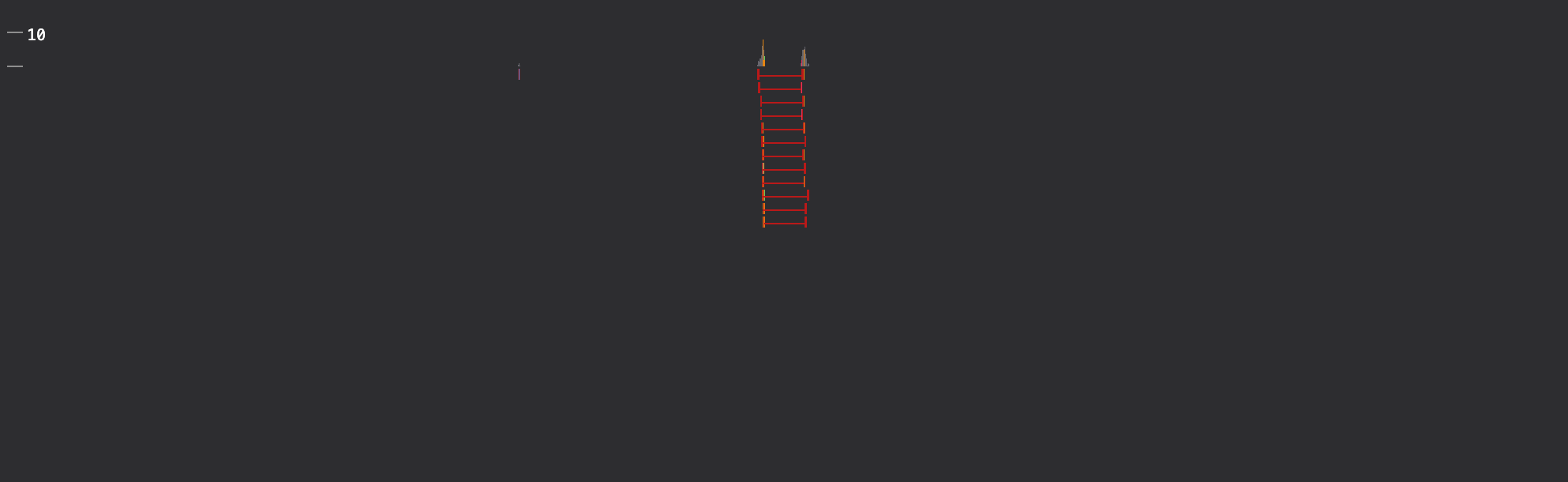

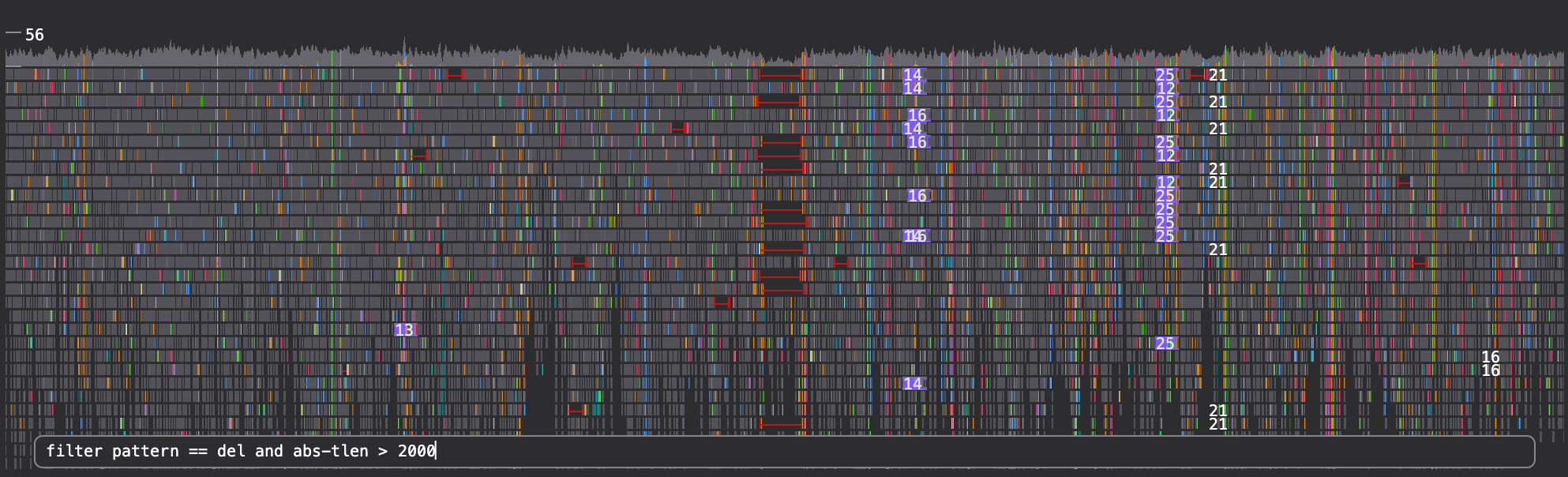

Reads with a deletion pattern and absolute insert-size > 2000

Reads mapping to a specific region

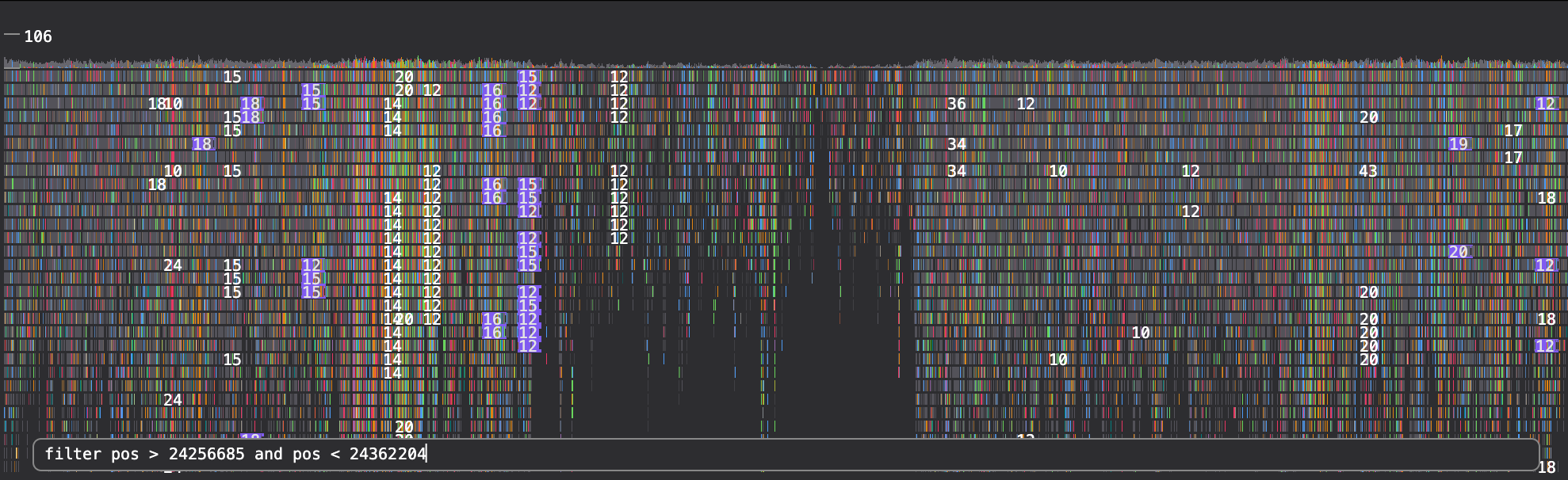

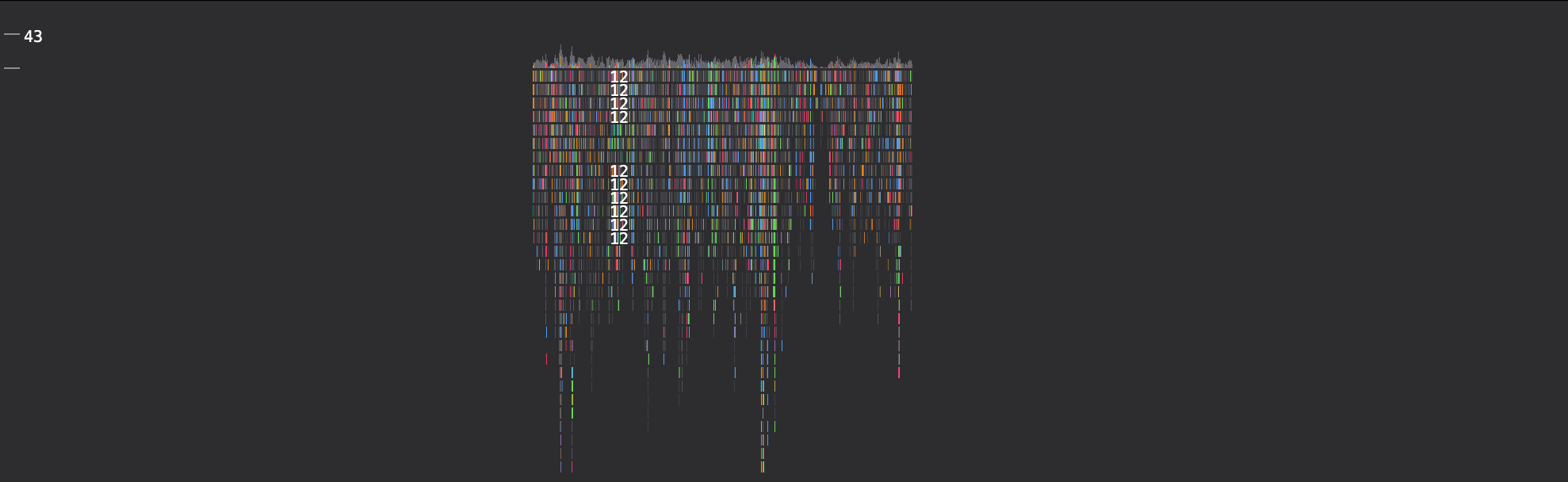

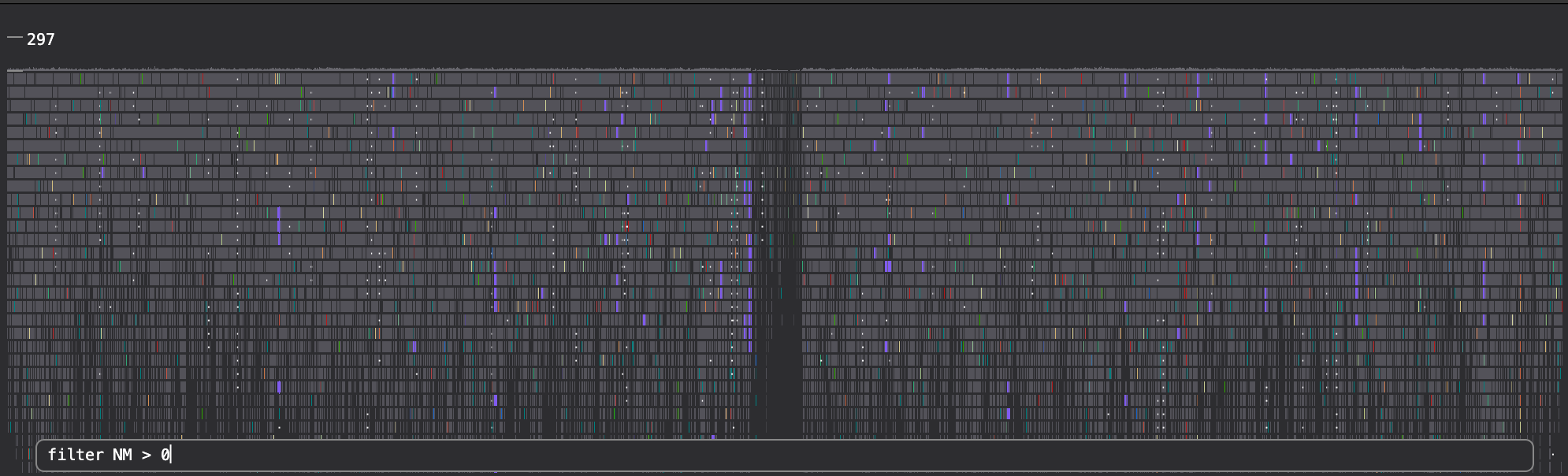

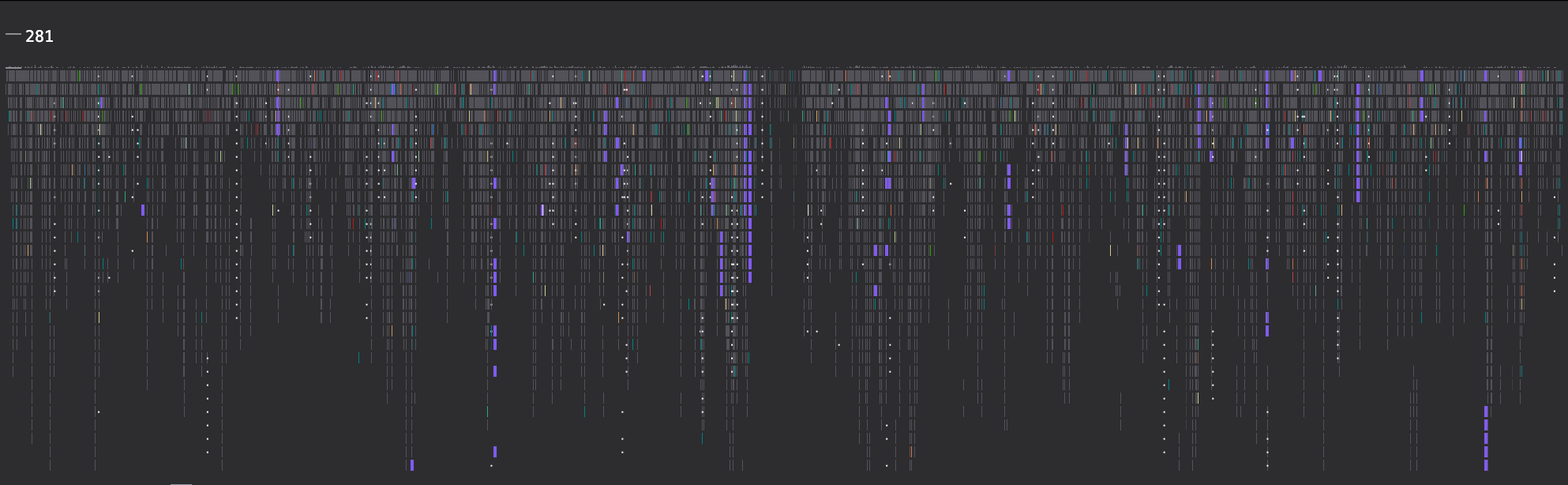

Reads with an edit distance > 0, using the NM sam tag

Reads with a kmer and cigar pattern

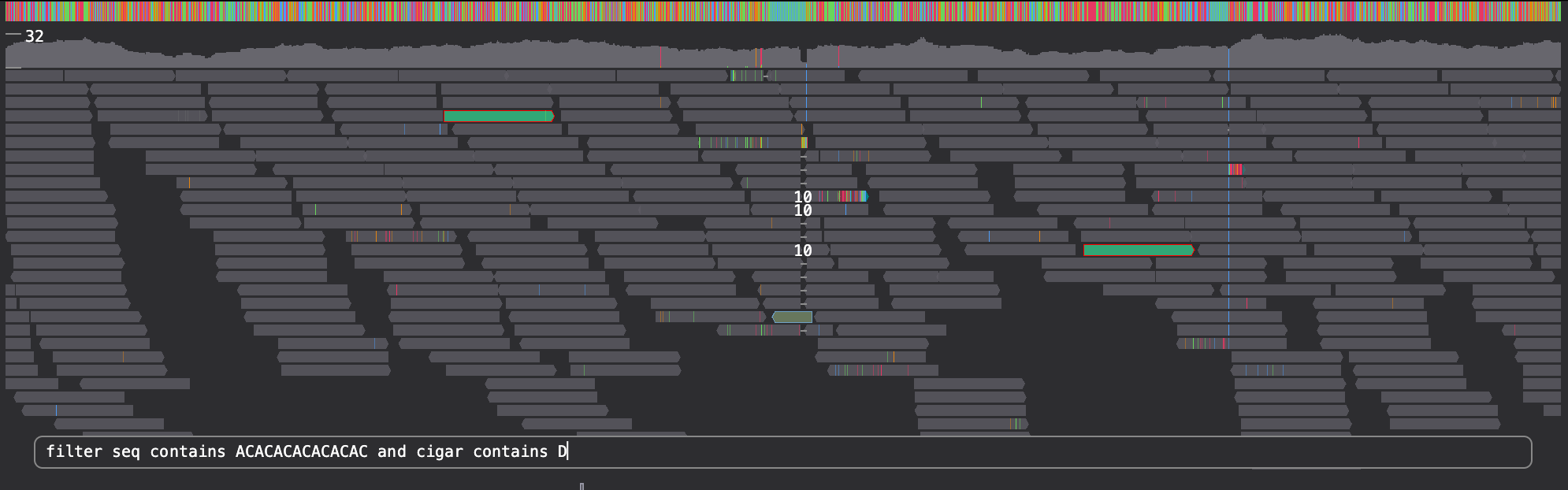

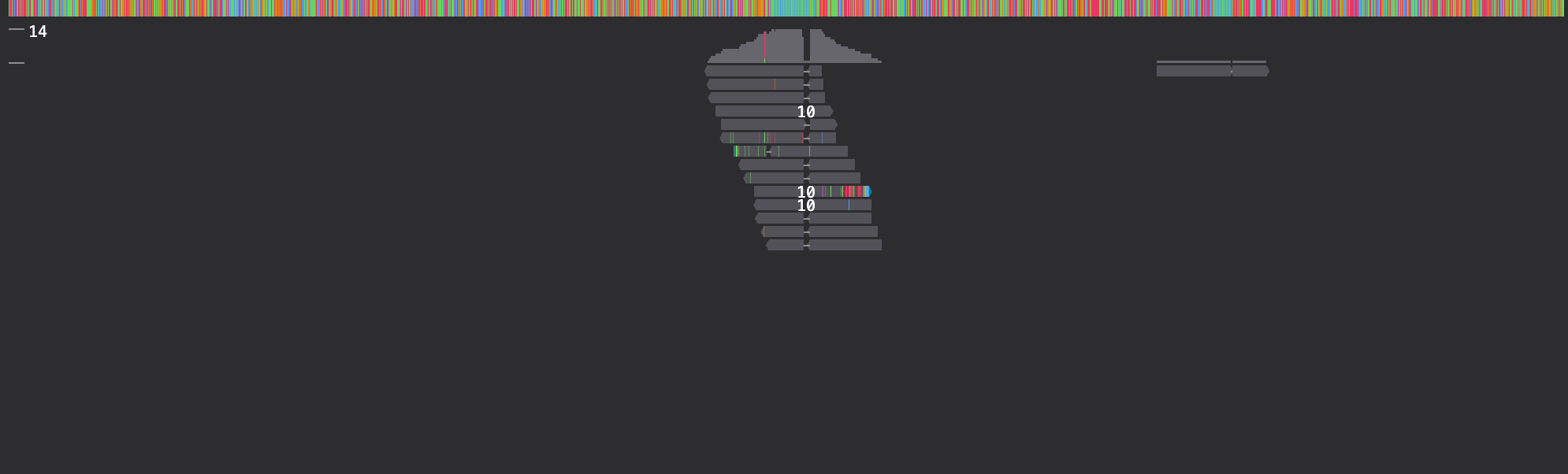

Reads with mate on chr7

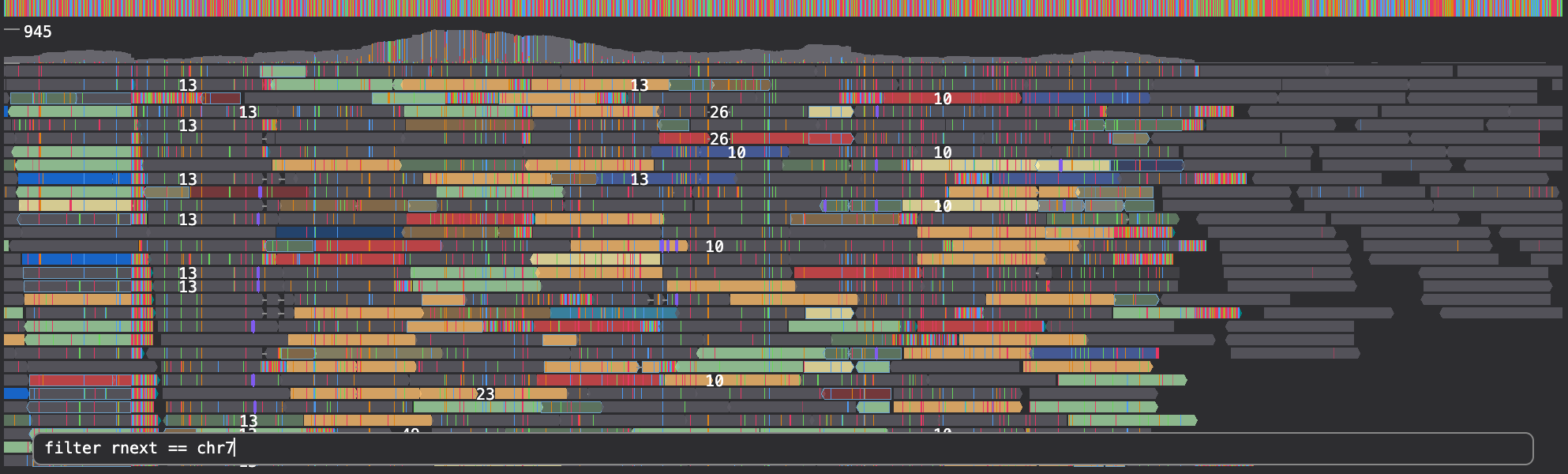

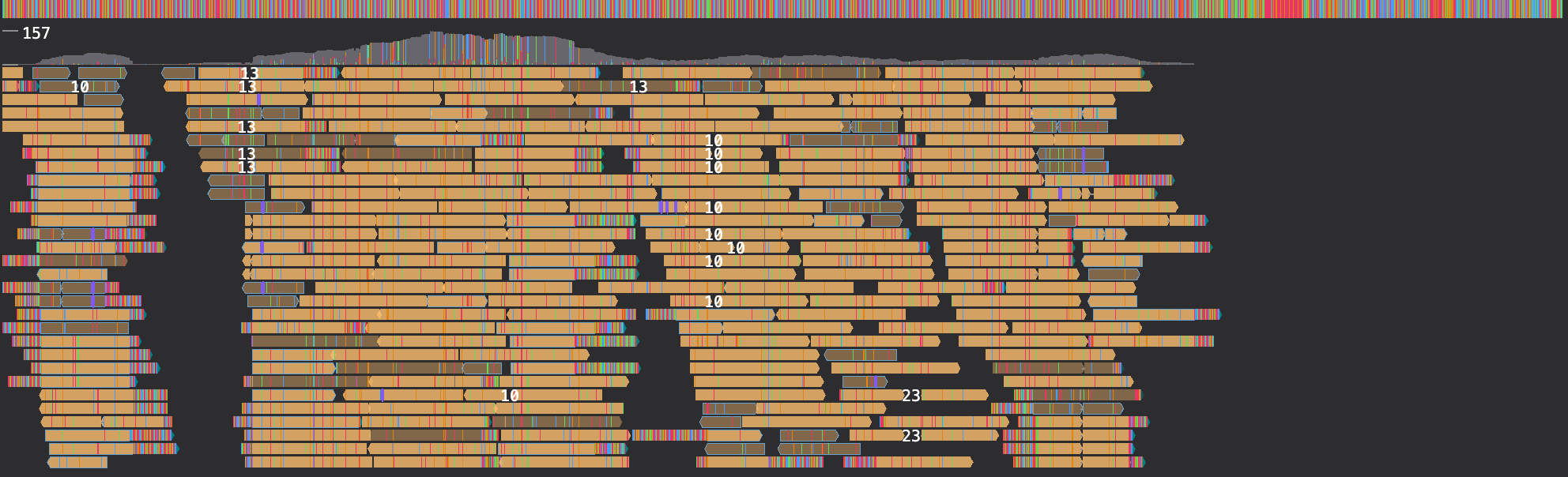

Keep high quality reads, no N in sequence, mapq >= 30 and edit-distance < 5 and no hard-clips

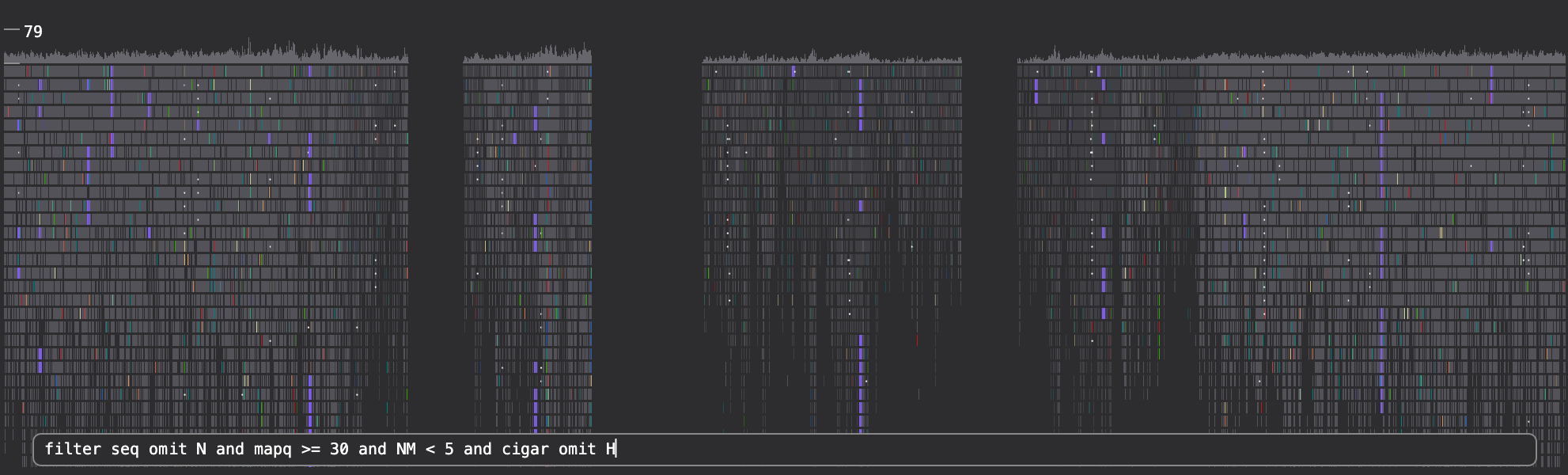

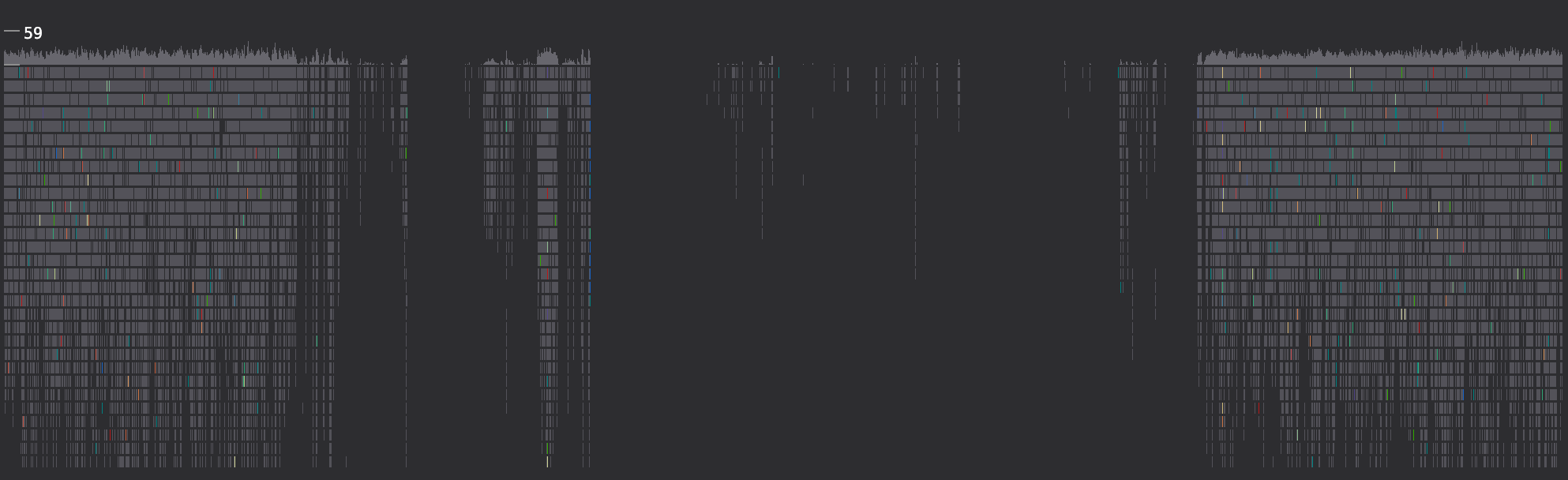

**Supplementary Figure 4. Example filtering commands.** A description of the filtering command is shown above the left-hand-size image, while the command itself is visible in the image. The result of the filtering command is shown on the right.

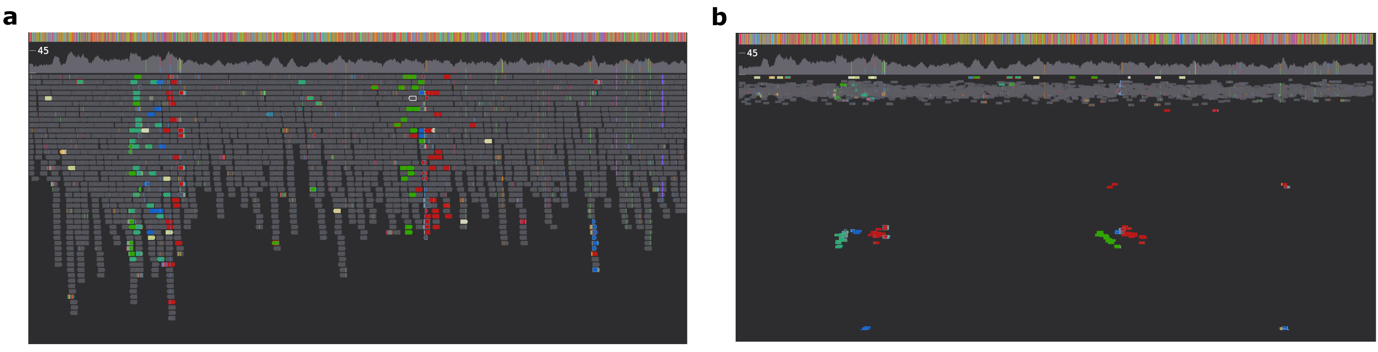

**Supplementary Figure 5. Scaling read position by template length.** The “TLEN” variable encoded in a bam file can be used to scale the y-position of reads using the “tlen-y’ command (a, b). This can be helpful when inspecting complex SV events using paired end reads.

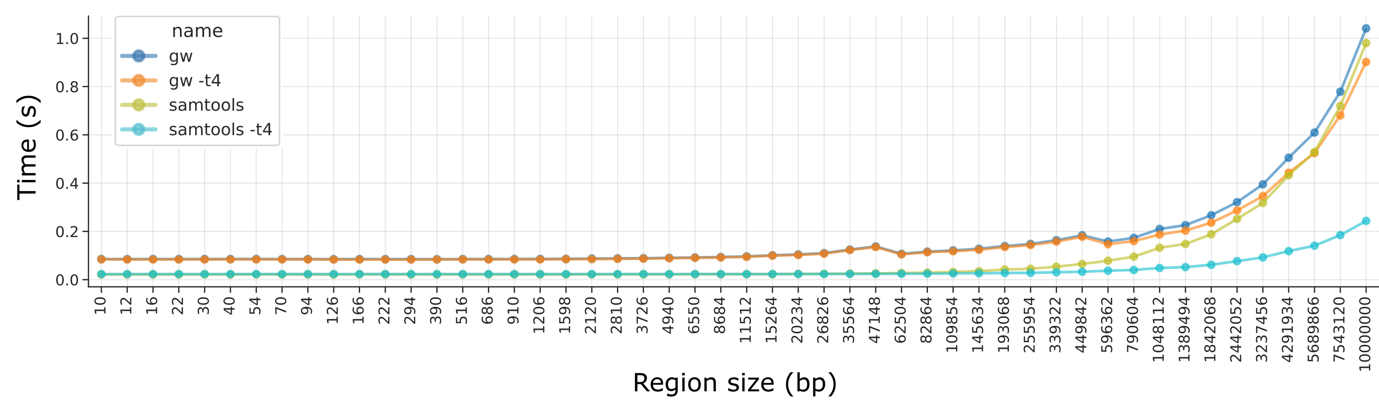

**Supplementary Figure 6. GW runtime vs ‘samtools view -c’.** Illumina reads at 40X coverage were used and the total execution time of GW was measured when drawing 20 random genome locations, including the time taken to save an image to disk. “-t4” indicates 4 threads were used.

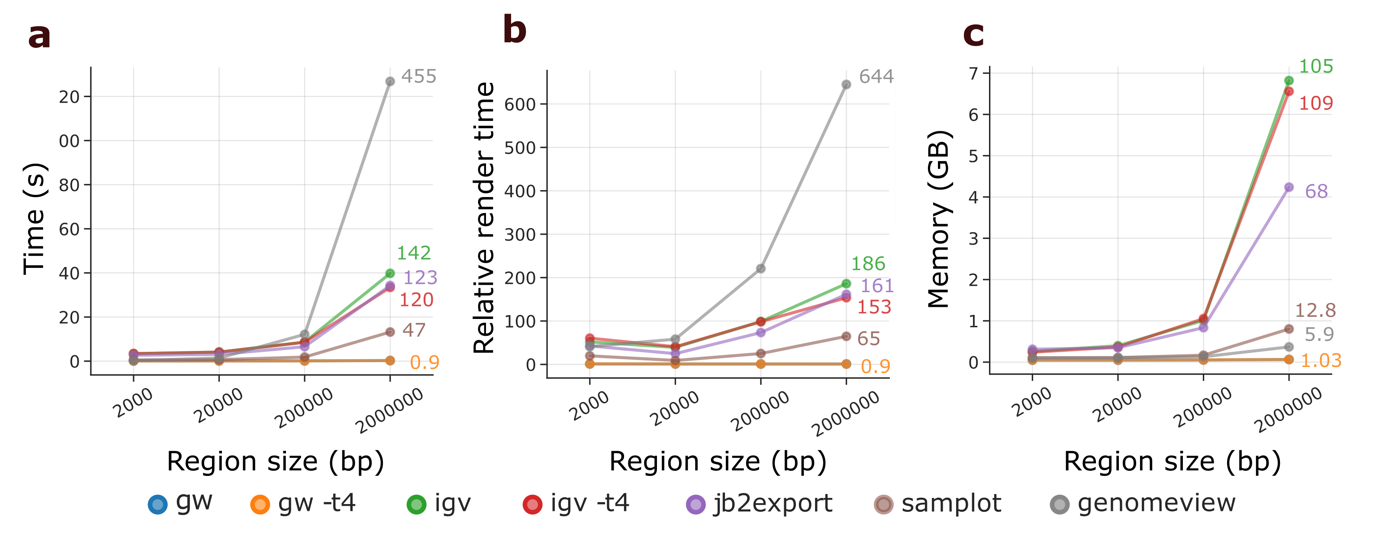

**Supplementary Figure 7. Illumina paired-end 40X benchmark.** Time (a), relative render time (b) and memory (c) were assessed when generating images of 20 random locations per region size. Colored numbers indicate the fold-difference of tools when compared to GW run in single-thread mode.

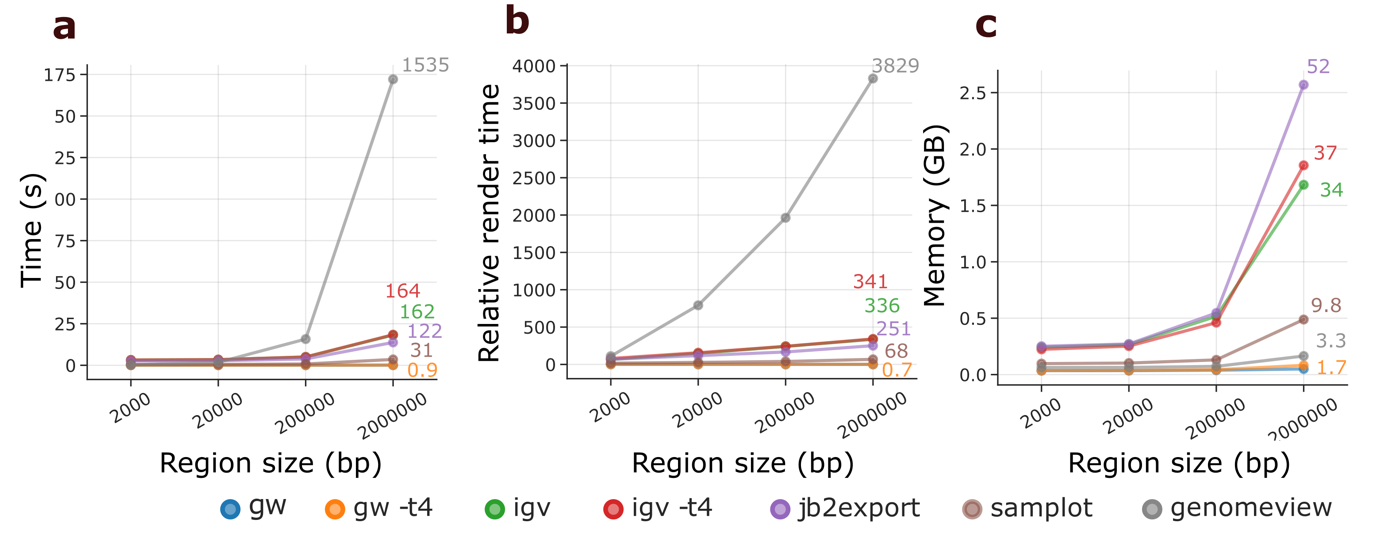

**Supplementary Figure 8. PacBio HiFi 8X read benchmark.**

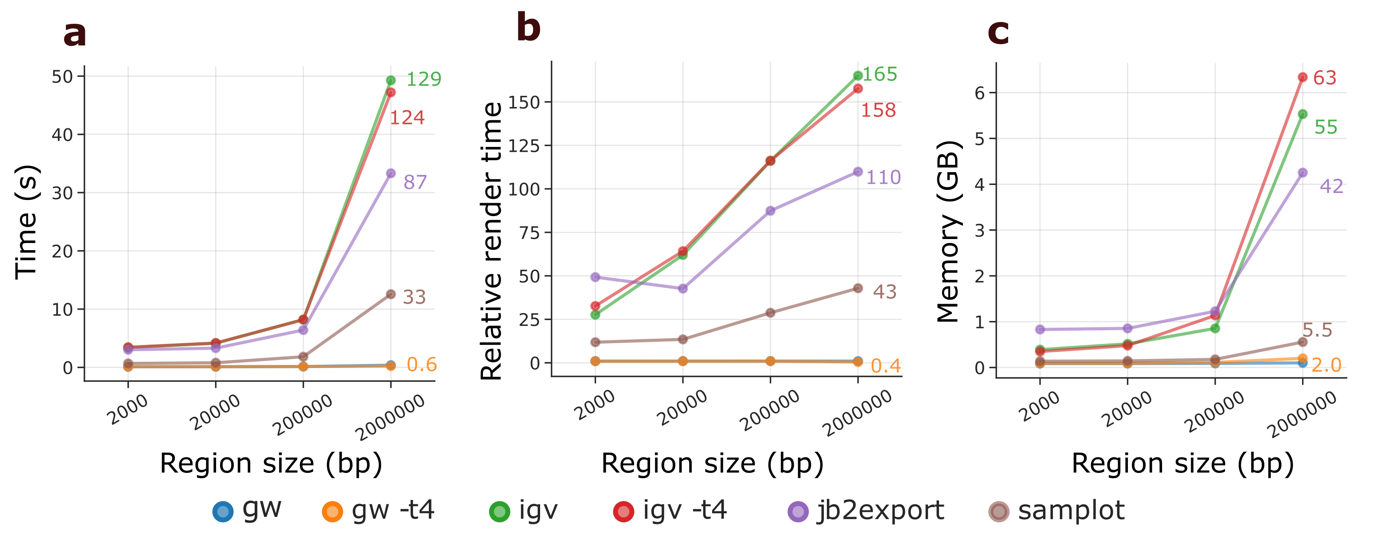

**Supplementary Figure 9. ONT Kit14 45X benchmark.**

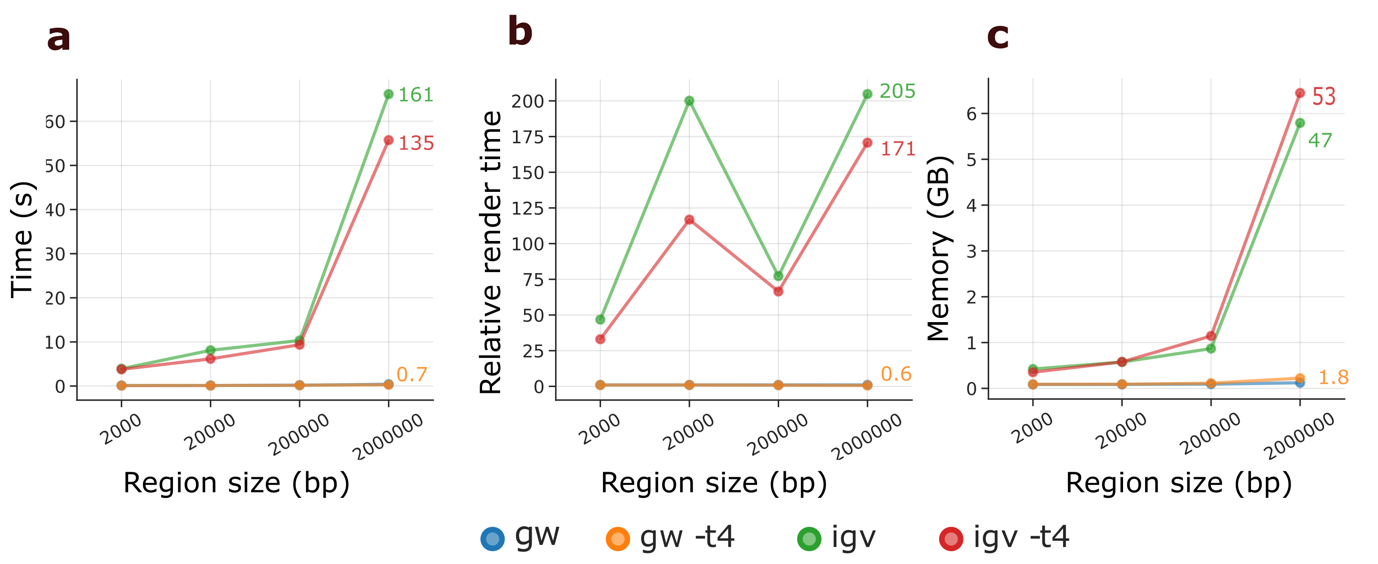

**Supplementary Figure 10. ONT Kit14 45X with modifications shown benchmark.**

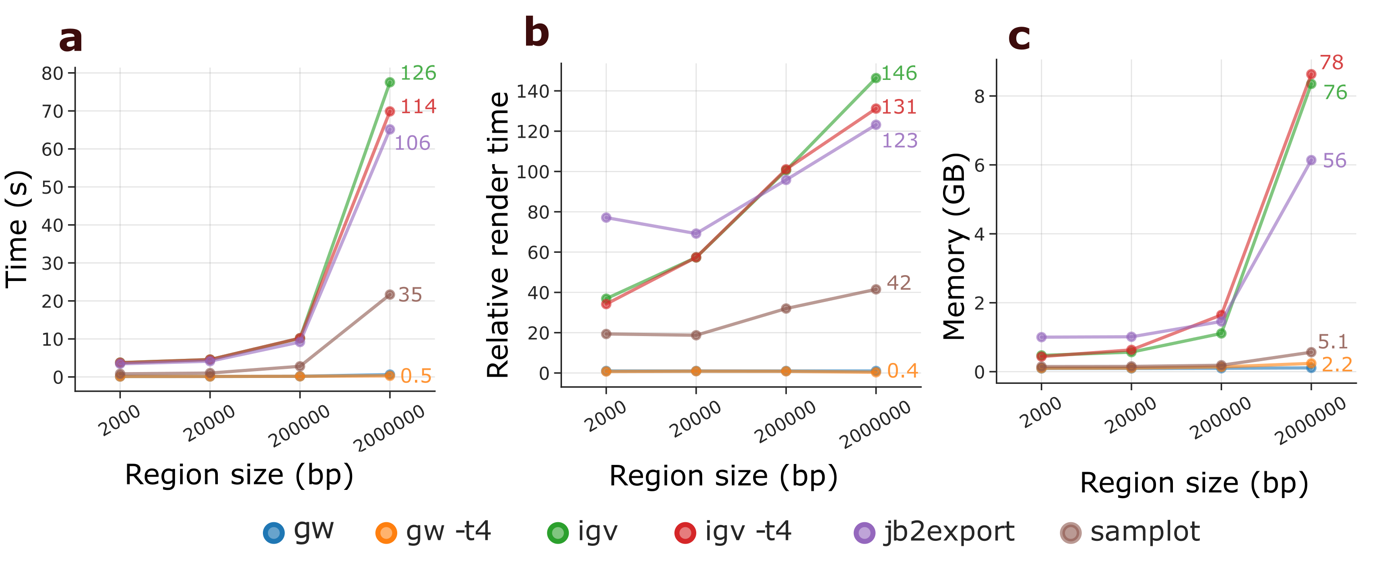

**Supplementary Figure 11. ONT Kit14 90X benchmark.**

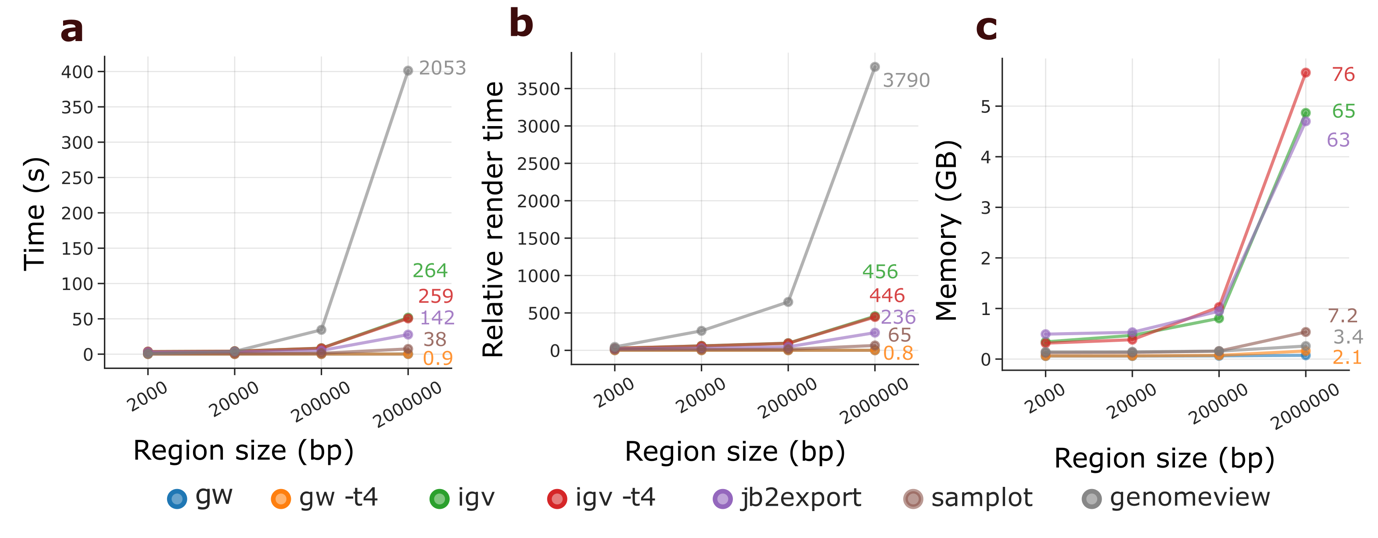

**Supplementary Figure 12. ONT R9.4.1 12.5X benchmark.**
