## Supplementary_Data for "GW: ultra-fast chromosome-scale visualisation of genomics data"

### Slide 1
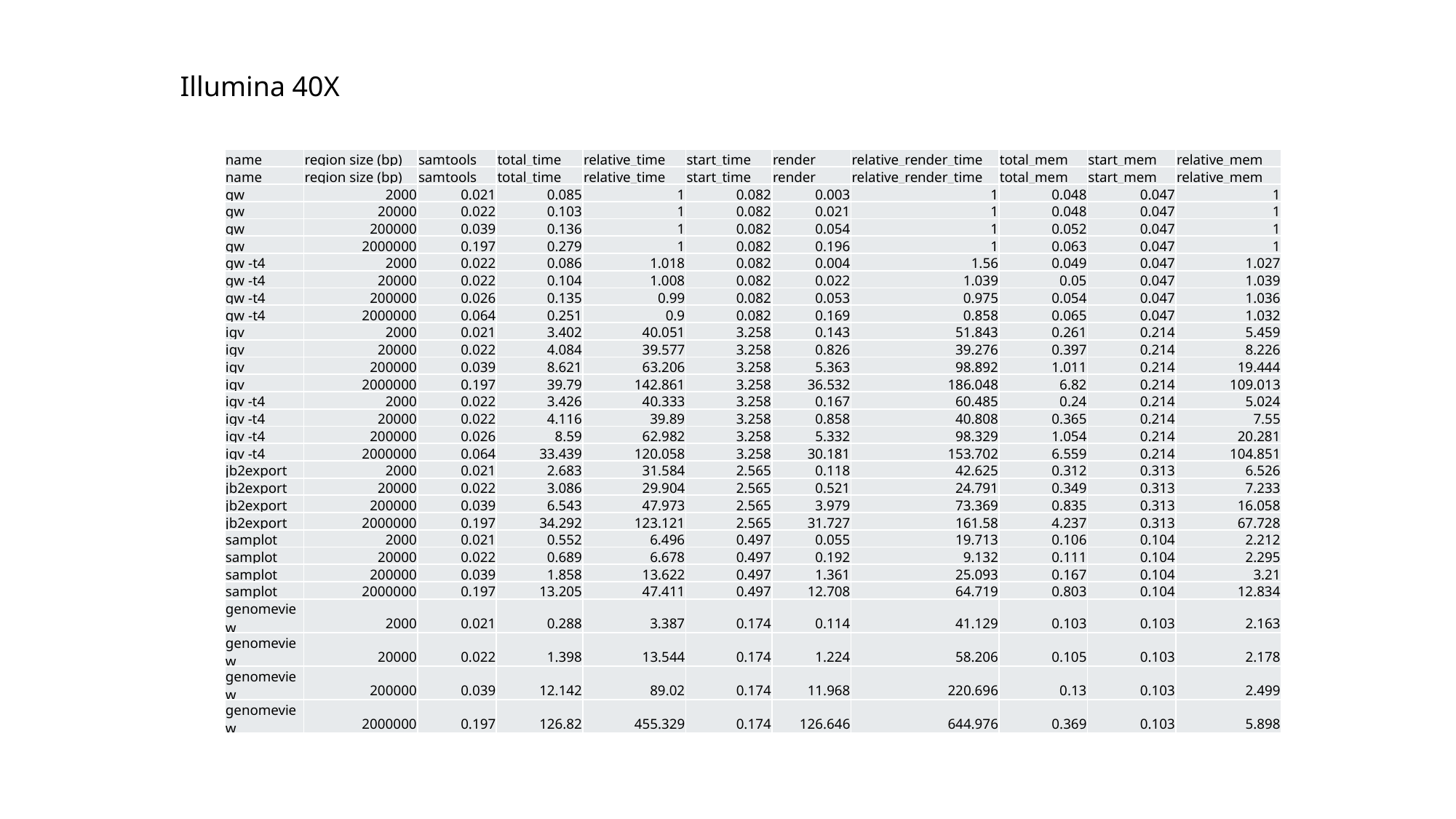

Illumina 40X
| name | region size (bp) | samtools | total\_time | relative\_time | start\_time | render | relative\_render\_time | total\_mem | start\_mem | relative\_mem |
| --- | --- | --- | --- | --- | --- | --- | --- | --- | --- | --- |
| name | region size (bp) | samtools | total\_time | relative\_time | start\_time | render | relative\_render\_time | total\_mem | start\_mem | relative\_mem |
| gw | 2000 | 0.021 | 0.085 | 1 | 0.082 | 0.003 | 1 | 0.048 | 0.047 | 1 |
| gw | 20000 | 0.022 | 0.103 | 1 | 0.082 | 0.021 | 1 | 0.048 | 0.047 | 1 |
| gw | 200000 | 0.039 | 0.136 | 1 | 0.082 | 0.054 | 1 | 0.052 | 0.047 | 1 |
| gw | 2000000 | 0.197 | 0.279 | 1 | 0.082 | 0.196 | 1 | 0.063 | 0.047 | 1 |
| gw -t4 | 2000 | 0.022 | 0.086 | 1.018 | 0.082 | 0.004 | 1.56 | 0.049 | 0.047 | 1.027 |
| gw -t4 | 20000 | 0.022 | 0.104 | 1.008 | 0.082 | 0.022 | 1.039 | 0.05 | 0.047 | 1.039 |
| gw -t4 | 200000 | 0.026 | 0.135 | 0.99 | 0.082 | 0.053 | 0.975 | 0.054 | 0.047 | 1.036 |
| gw -t4 | 2000000 | 0.064 | 0.251 | 0.9 | 0.082 | 0.169 | 0.858 | 0.065 | 0.047 | 1.032 |
| igv | 2000 | 0.021 | 3.402 | 40.051 | 3.258 | 0.143 | 51.843 | 0.261 | 0.214 | 5.459 |
| igv | 20000 | 0.022 | 4.084 | 39.577 | 3.258 | 0.826 | 39.276 | 0.397 | 0.214 | 8.226 |
| igv | 200000 | 0.039 | 8.621 | 63.206 | 3.258 | 5.363 | 98.892 | 1.011 | 0.214 | 19.444 |
| igv | 2000000 | 0.197 | 39.79 | 142.861 | 3.258 | 36.532 | 186.048 | 6.82 | 0.214 | 109.013 |
| igv -t4 | 2000 | 0.022 | 3.426 | 40.333 | 3.258 | 0.167 | 60.485 | 0.24 | 0.214 | 5.024 |
| igv -t4 | 20000 | 0.022 | 4.116 | 39.89 | 3.258 | 0.858 | 40.808 | 0.365 | 0.214 | 7.55 |
| igv -t4 | 200000 | 0.026 | 8.59 | 62.982 | 3.258 | 5.332 | 98.329 | 1.054 | 0.214 | 20.281 |
| igv -t4 | 2000000 | 0.064 | 33.439 | 120.058 | 3.258 | 30.181 | 153.702 | 6.559 | 0.214 | 104.851 |
| jb2export | 2000 | 0.021 | 2.683 | 31.584 | 2.565 | 0.118 | 42.625 | 0.312 | 0.313 | 6.526 |
| jb2export | 20000 | 0.022 | 3.086 | 29.904 | 2.565 | 0.521 | 24.791 | 0.349 | 0.313 | 7.233 |
| jb2export | 200000 | 0.039 | 6.543 | 47.973 | 2.565 | 3.979 | 73.369 | 0.835 | 0.313 | 16.058 |
| jb2export | 2000000 | 0.197 | 34.292 | 123.121 | 2.565 | 31.727 | 161.58 | 4.237 | 0.313 | 67.728 |
| samplot | 2000 | 0.021 | 0.552 | 6.496 | 0.497 | 0.055 | 19.713 | 0.106 | 0.104 | 2.212 |
| samplot | 20000 | 0.022 | 0.689 | 6.678 | 0.497 | 0.192 | 9.132 | 0.111 | 0.104 | 2.295 |
| samplot | 200000 | 0.039 | 1.858 | 13.622 | 0.497 | 1.361 | 25.093 | 0.167 | 0.104 | 3.21 |
| samplot | 2000000 | 0.197 | 13.205 | 47.411 | 0.497 | 12.708 | 64.719 | 0.803 | 0.104 | 12.834 |
| genomeview | 2000 | 0.021 | 0.288 | 3.387 | 0.174 | 0.114 | 41.129 | 0.103 | 0.103 | 2.163 |
| genomeview | 20000 | 0.022 | 1.398 | 13.544 | 0.174 | 1.224 | 58.206 | 0.105 | 0.103 | 2.178 |
| genomeview | 200000 | 0.039 | 12.142 | 89.02 | 0.174 | 11.968 | 220.696 | 0.13 | 0.103 | 2.499 |
| genomeview | 2000000 | 0.197 | 126.82 | 455.329 | 0.174 | 126.646 | 644.976 | 0.369 | 0.103 | 5.898 |

### Slide 2
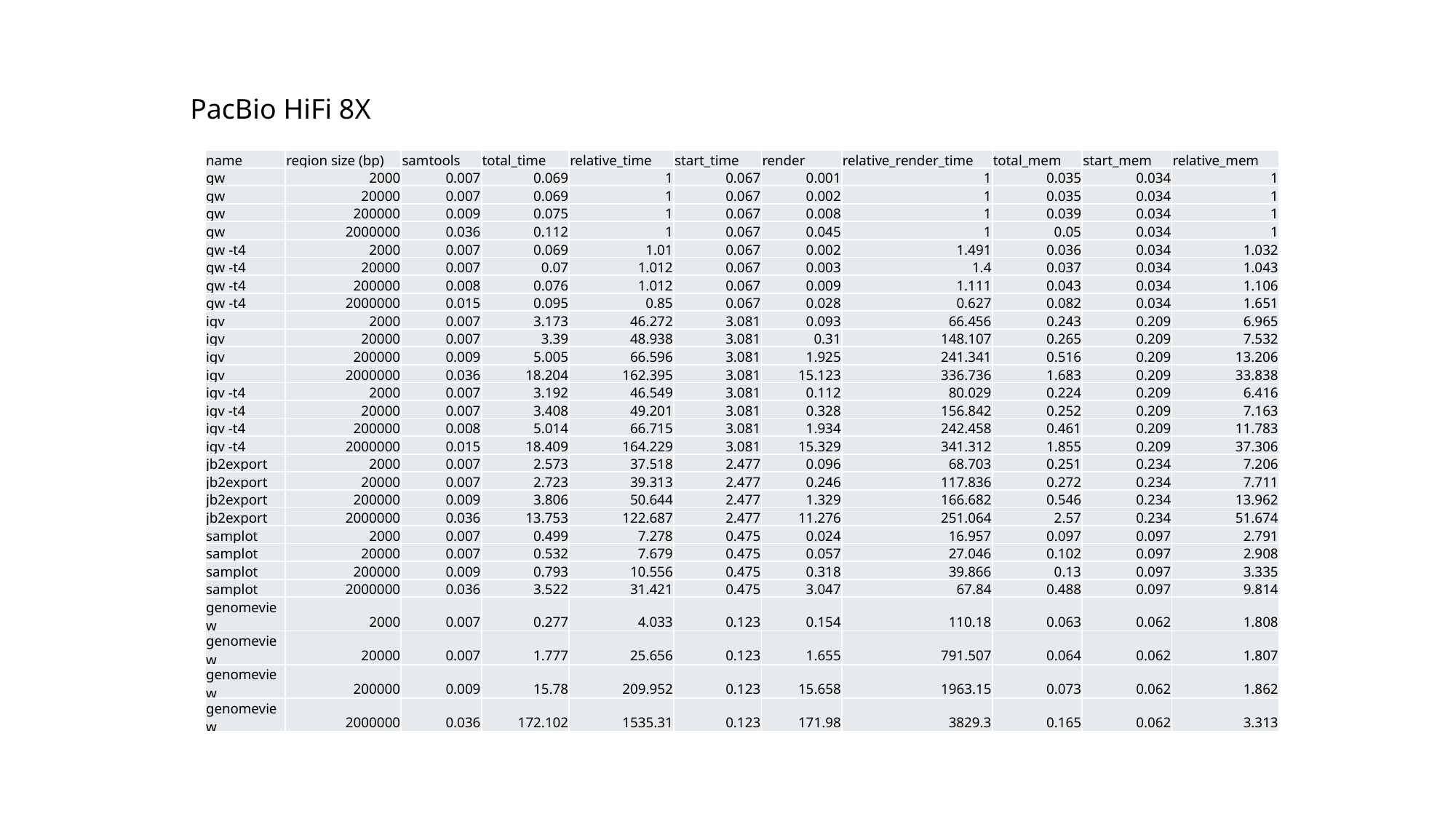

PacBio HiFi 8X
| name | region size (bp) | samtools | total\_time | relative\_time | start\_time | render | relative\_render\_time | total\_mem | start\_mem | relative\_mem |
| --- | --- | --- | --- | --- | --- | --- | --- | --- | --- | --- |
| gw | 2000 | 0.007 | 0.069 | 1 | 0.067 | 0.001 | 1 | 0.035 | 0.034 | 1 |
| gw | 20000 | 0.007 | 0.069 | 1 | 0.067 | 0.002 | 1 | 0.035 | 0.034 | 1 |
| gw | 200000 | 0.009 | 0.075 | 1 | 0.067 | 0.008 | 1 | 0.039 | 0.034 | 1 |
| gw | 2000000 | 0.036 | 0.112 | 1 | 0.067 | 0.045 | 1 | 0.05 | 0.034 | 1 |
| gw -t4 | 2000 | 0.007 | 0.069 | 1.01 | 0.067 | 0.002 | 1.491 | 0.036 | 0.034 | 1.032 |
| gw -t4 | 20000 | 0.007 | 0.07 | 1.012 | 0.067 | 0.003 | 1.4 | 0.037 | 0.034 | 1.043 |
| gw -t4 | 200000 | 0.008 | 0.076 | 1.012 | 0.067 | 0.009 | 1.111 | 0.043 | 0.034 | 1.106 |
| gw -t4 | 2000000 | 0.015 | 0.095 | 0.85 | 0.067 | 0.028 | 0.627 | 0.082 | 0.034 | 1.651 |
| igv | 2000 | 0.007 | 3.173 | 46.272 | 3.081 | 0.093 | 66.456 | 0.243 | 0.209 | 6.965 |
| igv | 20000 | 0.007 | 3.39 | 48.938 | 3.081 | 0.31 | 148.107 | 0.265 | 0.209 | 7.532 |
| igv | 200000 | 0.009 | 5.005 | 66.596 | 3.081 | 1.925 | 241.341 | 0.516 | 0.209 | 13.206 |
| igv | 2000000 | 0.036 | 18.204 | 162.395 | 3.081 | 15.123 | 336.736 | 1.683 | 0.209 | 33.838 |
| igv -t4 | 2000 | 0.007 | 3.192 | 46.549 | 3.081 | 0.112 | 80.029 | 0.224 | 0.209 | 6.416 |
| igv -t4 | 20000 | 0.007 | 3.408 | 49.201 | 3.081 | 0.328 | 156.842 | 0.252 | 0.209 | 7.163 |
| igv -t4 | 200000 | 0.008 | 5.014 | 66.715 | 3.081 | 1.934 | 242.458 | 0.461 | 0.209 | 11.783 |
| igv -t4 | 2000000 | 0.015 | 18.409 | 164.229 | 3.081 | 15.329 | 341.312 | 1.855 | 0.209 | 37.306 |
| jb2export | 2000 | 0.007 | 2.573 | 37.518 | 2.477 | 0.096 | 68.703 | 0.251 | 0.234 | 7.206 |
| jb2export | 20000 | 0.007 | 2.723 | 39.313 | 2.477 | 0.246 | 117.836 | 0.272 | 0.234 | 7.711 |
| jb2export | 200000 | 0.009 | 3.806 | 50.644 | 2.477 | 1.329 | 166.682 | 0.546 | 0.234 | 13.962 |
| jb2export | 2000000 | 0.036 | 13.753 | 122.687 | 2.477 | 11.276 | 251.064 | 2.57 | 0.234 | 51.674 |
| samplot | 2000 | 0.007 | 0.499 | 7.278 | 0.475 | 0.024 | 16.957 | 0.097 | 0.097 | 2.791 |
| samplot | 20000 | 0.007 | 0.532 | 7.679 | 0.475 | 0.057 | 27.046 | 0.102 | 0.097 | 2.908 |
| samplot | 200000 | 0.009 | 0.793 | 10.556 | 0.475 | 0.318 | 39.866 | 0.13 | 0.097 | 3.335 |
| samplot | 2000000 | 0.036 | 3.522 | 31.421 | 0.475 | 3.047 | 67.84 | 0.488 | 0.097 | 9.814 |
| genomeview | 2000 | 0.007 | 0.277 | 4.033 | 0.123 | 0.154 | 110.18 | 0.063 | 0.062 | 1.808 |
| genomeview | 20000 | 0.007 | 1.777 | 25.656 | 0.123 | 1.655 | 791.507 | 0.064 | 0.062 | 1.807 |
| genomeview | 200000 | 0.009 | 15.78 | 209.952 | 0.123 | 15.658 | 1963.15 | 0.073 | 0.062 | 1.862 |
| genomeview | 2000000 | 0.036 | 172.102 | 1535.31 | 0.123 | 171.98 | 3829.3 | 0.165 | 0.062 | 3.313 |

### Slide 3
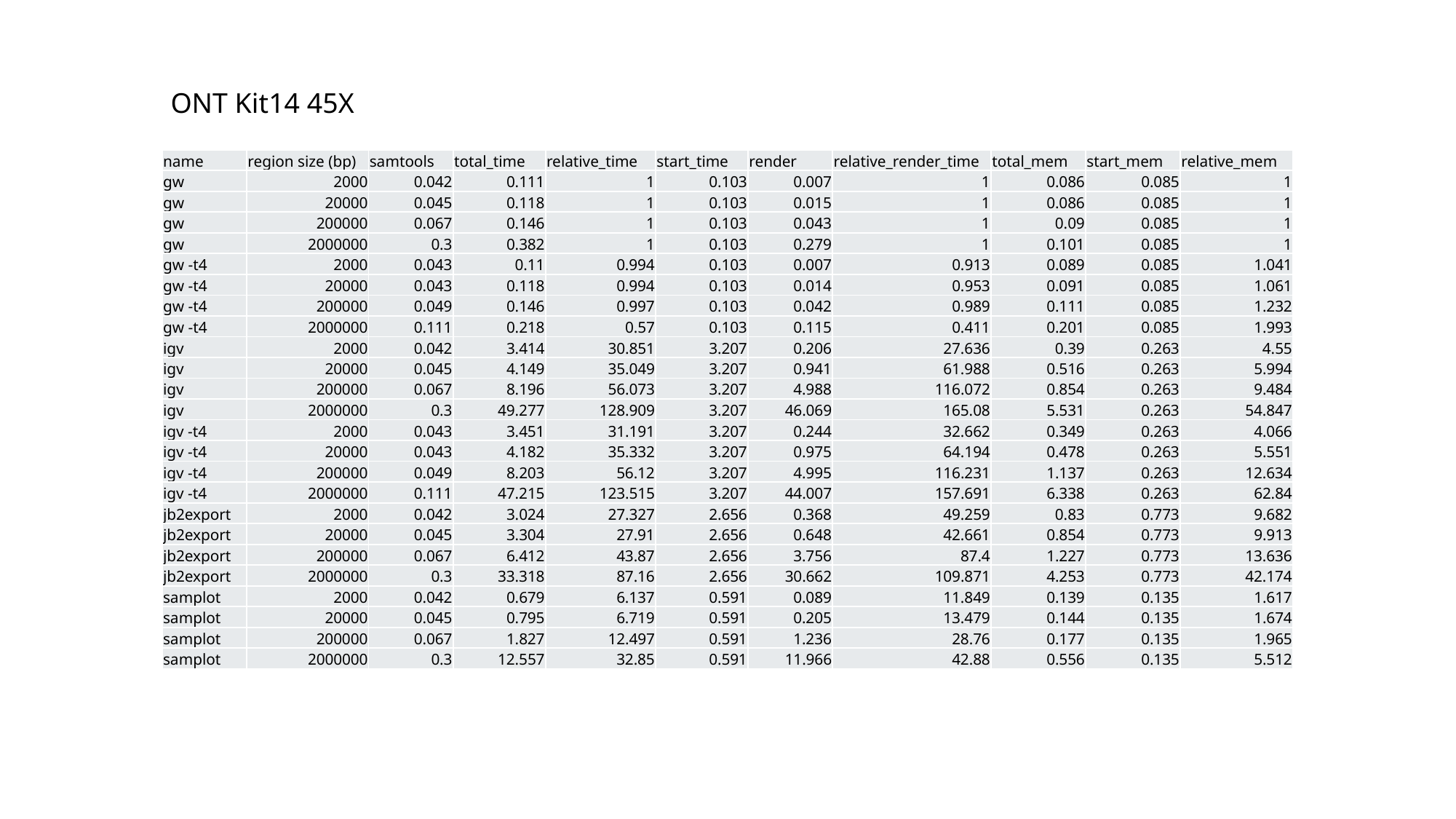

ONT Kit14 45X
| name | region size (bp) | samtools | total\_time | relative\_time | start\_time | render | relative\_render\_time | total\_mem | start\_mem | relative\_mem |
| --- | --- | --- | --- | --- | --- | --- | --- | --- | --- | --- |
| gw | 2000 | 0.042 | 0.111 | 1 | 0.103 | 0.007 | 1 | 0.086 | 0.085 | 1 |
| gw | 20000 | 0.045 | 0.118 | 1 | 0.103 | 0.015 | 1 | 0.086 | 0.085 | 1 |
| gw | 200000 | 0.067 | 0.146 | 1 | 0.103 | 0.043 | 1 | 0.09 | 0.085 | 1 |
| gw | 2000000 | 0.3 | 0.382 | 1 | 0.103 | 0.279 | 1 | 0.101 | 0.085 | 1 |
| gw -t4 | 2000 | 0.043 | 0.11 | 0.994 | 0.103 | 0.007 | 0.913 | 0.089 | 0.085 | 1.041 |
| gw -t4 | 20000 | 0.043 | 0.118 | 0.994 | 0.103 | 0.014 | 0.953 | 0.091 | 0.085 | 1.061 |
| gw -t4 | 200000 | 0.049 | 0.146 | 0.997 | 0.103 | 0.042 | 0.989 | 0.111 | 0.085 | 1.232 |
| gw -t4 | 2000000 | 0.111 | 0.218 | 0.57 | 0.103 | 0.115 | 0.411 | 0.201 | 0.085 | 1.993 |
| igv | 2000 | 0.042 | 3.414 | 30.851 | 3.207 | 0.206 | 27.636 | 0.39 | 0.263 | 4.55 |
| igv | 20000 | 0.045 | 4.149 | 35.049 | 3.207 | 0.941 | 61.988 | 0.516 | 0.263 | 5.994 |
| igv | 200000 | 0.067 | 8.196 | 56.073 | 3.207 | 4.988 | 116.072 | 0.854 | 0.263 | 9.484 |
| igv | 2000000 | 0.3 | 49.277 | 128.909 | 3.207 | 46.069 | 165.08 | 5.531 | 0.263 | 54.847 |
| igv -t4 | 2000 | 0.043 | 3.451 | 31.191 | 3.207 | 0.244 | 32.662 | 0.349 | 0.263 | 4.066 |
| igv -t4 | 20000 | 0.043 | 4.182 | 35.332 | 3.207 | 0.975 | 64.194 | 0.478 | 0.263 | 5.551 |
| igv -t4 | 200000 | 0.049 | 8.203 | 56.12 | 3.207 | 4.995 | 116.231 | 1.137 | 0.263 | 12.634 |
| igv -t4 | 2000000 | 0.111 | 47.215 | 123.515 | 3.207 | 44.007 | 157.691 | 6.338 | 0.263 | 62.84 |
| jb2export | 2000 | 0.042 | 3.024 | 27.327 | 2.656 | 0.368 | 49.259 | 0.83 | 0.773 | 9.682 |
| jb2export | 20000 | 0.045 | 3.304 | 27.91 | 2.656 | 0.648 | 42.661 | 0.854 | 0.773 | 9.913 |
| jb2export | 200000 | 0.067 | 6.412 | 43.87 | 2.656 | 3.756 | 87.4 | 1.227 | 0.773 | 13.636 |
| jb2export | 2000000 | 0.3 | 33.318 | 87.16 | 2.656 | 30.662 | 109.871 | 4.253 | 0.773 | 42.174 |
| samplot | 2000 | 0.042 | 0.679 | 6.137 | 0.591 | 0.089 | 11.849 | 0.139 | 0.135 | 1.617 |
| samplot | 20000 | 0.045 | 0.795 | 6.719 | 0.591 | 0.205 | 13.479 | 0.144 | 0.135 | 1.674 |
| samplot | 200000 | 0.067 | 1.827 | 12.497 | 0.591 | 1.236 | 28.76 | 0.177 | 0.135 | 1.965 |
| samplot | 2000000 | 0.3 | 12.557 | 32.85 | 0.591 | 11.966 | 42.88 | 0.556 | 0.135 | 5.512 |

### Slide 4
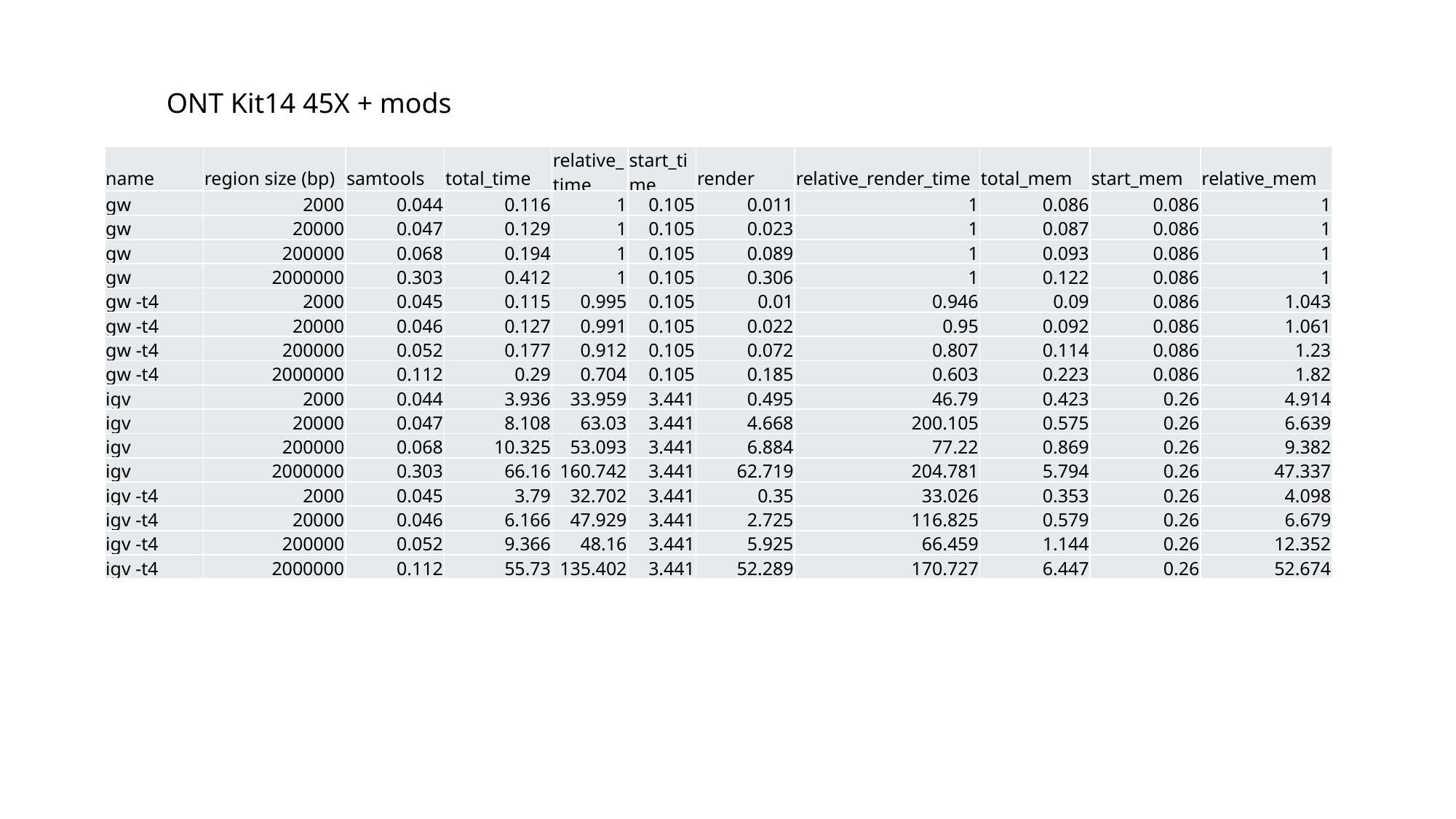

ONT Kit14 45X + mods
| name | region size (bp) | samtools | total\_time | relative\_time | start\_time | render | relative\_render\_time | total\_mem | start\_mem | relative\_mem |
| --- | --- | --- | --- | --- | --- | --- | --- | --- | --- | --- |
| gw | 2000 | 0.044 | 0.116 | 1 | 0.105 | 0.011 | 1 | 0.086 | 0.086 | 1 |
| gw | 20000 | 0.047 | 0.129 | 1 | 0.105 | 0.023 | 1 | 0.087 | 0.086 | 1 |
| gw | 200000 | 0.068 | 0.194 | 1 | 0.105 | 0.089 | 1 | 0.093 | 0.086 | 1 |
| gw | 2000000 | 0.303 | 0.412 | 1 | 0.105 | 0.306 | 1 | 0.122 | 0.086 | 1 |
| gw -t4 | 2000 | 0.045 | 0.115 | 0.995 | 0.105 | 0.01 | 0.946 | 0.09 | 0.086 | 1.043 |
| gw -t4 | 20000 | 0.046 | 0.127 | 0.991 | 0.105 | 0.022 | 0.95 | 0.092 | 0.086 | 1.061 |
| gw -t4 | 200000 | 0.052 | 0.177 | 0.912 | 0.105 | 0.072 | 0.807 | 0.114 | 0.086 | 1.23 |
| gw -t4 | 2000000 | 0.112 | 0.29 | 0.704 | 0.105 | 0.185 | 0.603 | 0.223 | 0.086 | 1.82 |
| igv | 2000 | 0.044 | 3.936 | 33.959 | 3.441 | 0.495 | 46.79 | 0.423 | 0.26 | 4.914 |
| igv | 20000 | 0.047 | 8.108 | 63.03 | 3.441 | 4.668 | 200.105 | 0.575 | 0.26 | 6.639 |
| igv | 200000 | 0.068 | 10.325 | 53.093 | 3.441 | 6.884 | 77.22 | 0.869 | 0.26 | 9.382 |
| igv | 2000000 | 0.303 | 66.16 | 160.742 | 3.441 | 62.719 | 204.781 | 5.794 | 0.26 | 47.337 |
| igv -t4 | 2000 | 0.045 | 3.79 | 32.702 | 3.441 | 0.35 | 33.026 | 0.353 | 0.26 | 4.098 |
| igv -t4 | 20000 | 0.046 | 6.166 | 47.929 | 3.441 | 2.725 | 116.825 | 0.579 | 0.26 | 6.679 |
| igv -t4 | 200000 | 0.052 | 9.366 | 48.16 | 3.441 | 5.925 | 66.459 | 1.144 | 0.26 | 12.352 |
| igv -t4 | 2000000 | 0.112 | 55.73 | 135.402 | 3.441 | 52.289 | 170.727 | 6.447 | 0.26 | 52.674 |

### Slide 5
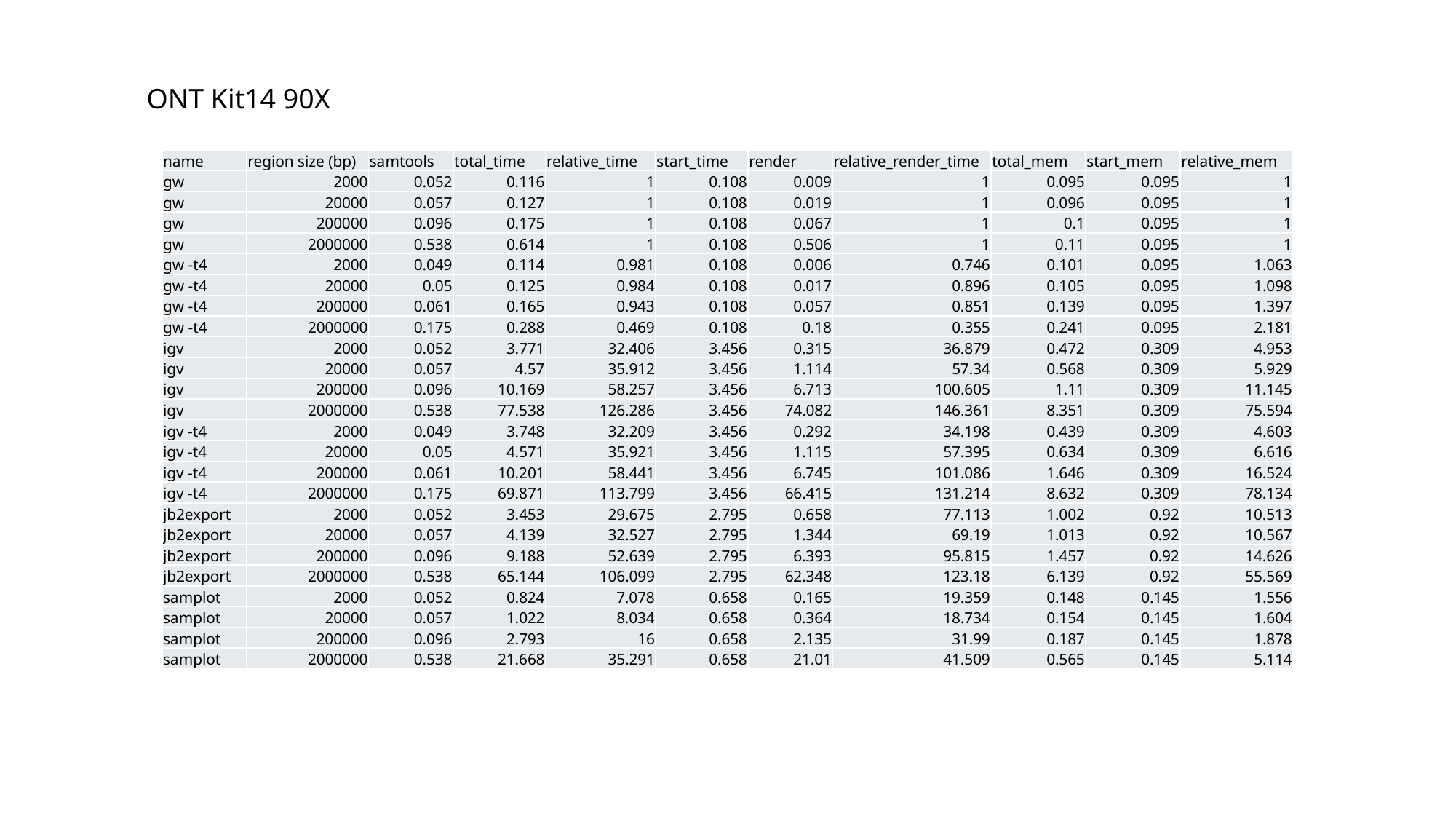

ONT Kit14 90X
| name | region size (bp) | samtools | total\_time | relative\_time | start\_time | render | relative\_render\_time | total\_mem | start\_mem | relative\_mem |
| --- | --- | --- | --- | --- | --- | --- | --- | --- | --- | --- |
| gw | 2000 | 0.052 | 0.116 | 1 | 0.108 | 0.009 | 1 | 0.095 | 0.095 | 1 |
| gw | 20000 | 0.057 | 0.127 | 1 | 0.108 | 0.019 | 1 | 0.096 | 0.095 | 1 |
| gw | 200000 | 0.096 | 0.175 | 1 | 0.108 | 0.067 | 1 | 0.1 | 0.095 | 1 |
| gw | 2000000 | 0.538 | 0.614 | 1 | 0.108 | 0.506 | 1 | 0.11 | 0.095 | 1 |
| gw -t4 | 2000 | 0.049 | 0.114 | 0.981 | 0.108 | 0.006 | 0.746 | 0.101 | 0.095 | 1.063 |
| gw -t4 | 20000 | 0.05 | 0.125 | 0.984 | 0.108 | 0.017 | 0.896 | 0.105 | 0.095 | 1.098 |
| gw -t4 | 200000 | 0.061 | 0.165 | 0.943 | 0.108 | 0.057 | 0.851 | 0.139 | 0.095 | 1.397 |
| gw -t4 | 2000000 | 0.175 | 0.288 | 0.469 | 0.108 | 0.18 | 0.355 | 0.241 | 0.095 | 2.181 |
| igv | 2000 | 0.052 | 3.771 | 32.406 | 3.456 | 0.315 | 36.879 | 0.472 | 0.309 | 4.953 |
| igv | 20000 | 0.057 | 4.57 | 35.912 | 3.456 | 1.114 | 57.34 | 0.568 | 0.309 | 5.929 |
| igv | 200000 | 0.096 | 10.169 | 58.257 | 3.456 | 6.713 | 100.605 | 1.11 | 0.309 | 11.145 |
| igv | 2000000 | 0.538 | 77.538 | 126.286 | 3.456 | 74.082 | 146.361 | 8.351 | 0.309 | 75.594 |
| igv -t4 | 2000 | 0.049 | 3.748 | 32.209 | 3.456 | 0.292 | 34.198 | 0.439 | 0.309 | 4.603 |
| igv -t4 | 20000 | 0.05 | 4.571 | 35.921 | 3.456 | 1.115 | 57.395 | 0.634 | 0.309 | 6.616 |
| igv -t4 | 200000 | 0.061 | 10.201 | 58.441 | 3.456 | 6.745 | 101.086 | 1.646 | 0.309 | 16.524 |
| igv -t4 | 2000000 | 0.175 | 69.871 | 113.799 | 3.456 | 66.415 | 131.214 | 8.632 | 0.309 | 78.134 |
| jb2export | 2000 | 0.052 | 3.453 | 29.675 | 2.795 | 0.658 | 77.113 | 1.002 | 0.92 | 10.513 |
| jb2export | 20000 | 0.057 | 4.139 | 32.527 | 2.795 | 1.344 | 69.19 | 1.013 | 0.92 | 10.567 |
| jb2export | 200000 | 0.096 | 9.188 | 52.639 | 2.795 | 6.393 | 95.815 | 1.457 | 0.92 | 14.626 |
| jb2export | 2000000 | 0.538 | 65.144 | 106.099 | 2.795 | 62.348 | 123.18 | 6.139 | 0.92 | 55.569 |
| samplot | 2000 | 0.052 | 0.824 | 7.078 | 0.658 | 0.165 | 19.359 | 0.148 | 0.145 | 1.556 |
| samplot | 20000 | 0.057 | 1.022 | 8.034 | 0.658 | 0.364 | 18.734 | 0.154 | 0.145 | 1.604 |
| samplot | 200000 | 0.096 | 2.793 | 16 | 0.658 | 2.135 | 31.99 | 0.187 | 0.145 | 1.878 |
| samplot | 2000000 | 0.538 | 21.668 | 35.291 | 0.658 | 21.01 | 41.509 | 0.565 | 0.145 | 5.114 |

### Slide 6
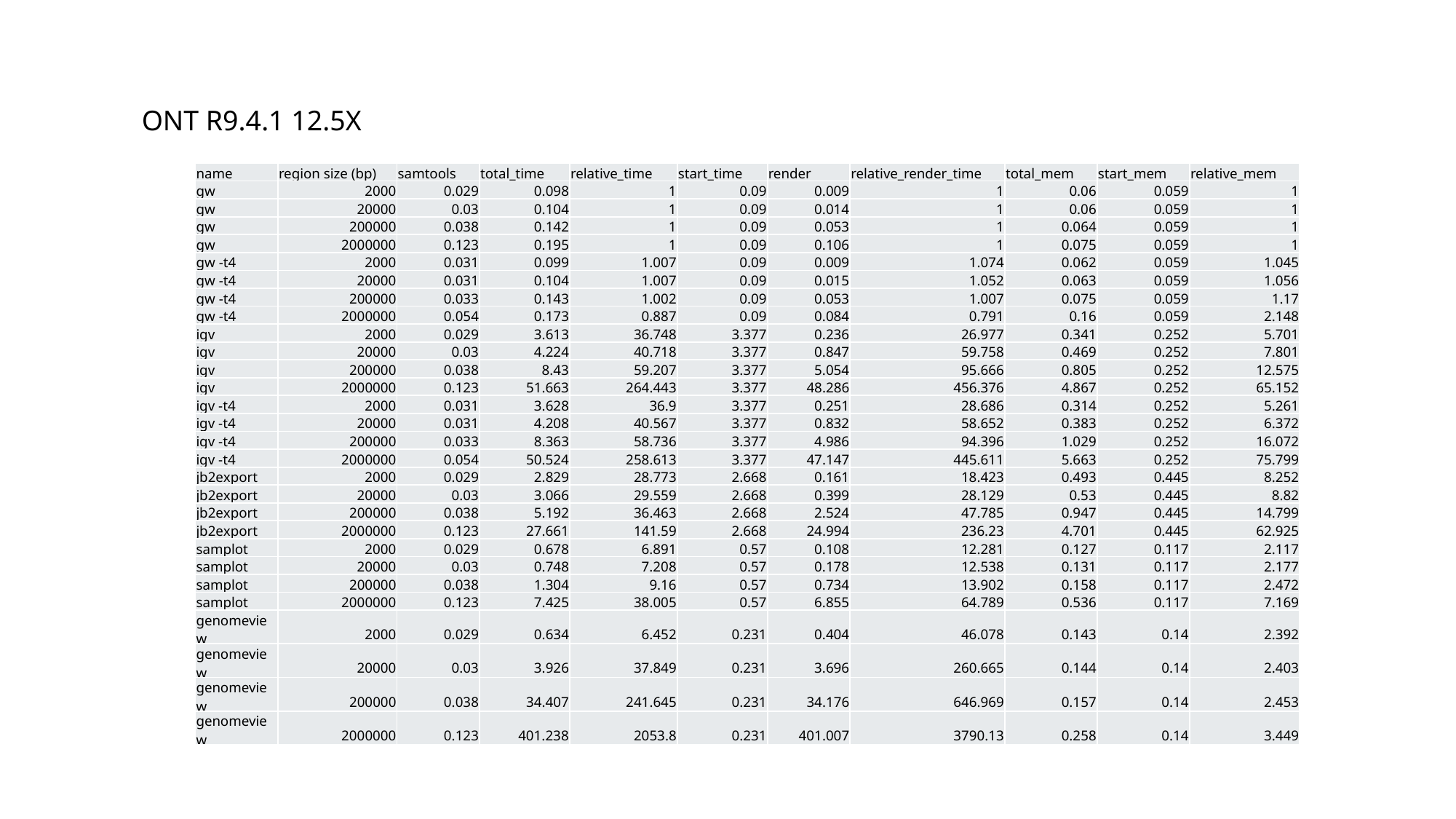

ONT R9.4.1 12.5X
| name | region size (bp) | samtools | total\_time | relative\_time | start\_time | render | relative\_render\_time | total\_mem | start\_mem | relative\_mem |
| --- | --- | --- | --- | --- | --- | --- | --- | --- | --- | --- |
| gw | 2000 | 0.029 | 0.098 | 1 | 0.09 | 0.009 | 1 | 0.06 | 0.059 | 1 |
| gw | 20000 | 0.03 | 0.104 | 1 | 0.09 | 0.014 | 1 | 0.06 | 0.059 | 1 |
| gw | 200000 | 0.038 | 0.142 | 1 | 0.09 | 0.053 | 1 | 0.064 | 0.059 | 1 |
| gw | 2000000 | 0.123 | 0.195 | 1 | 0.09 | 0.106 | 1 | 0.075 | 0.059 | 1 |
| gw -t4 | 2000 | 0.031 | 0.099 | 1.007 | 0.09 | 0.009 | 1.074 | 0.062 | 0.059 | 1.045 |
| gw -t4 | 20000 | 0.031 | 0.104 | 1.007 | 0.09 | 0.015 | 1.052 | 0.063 | 0.059 | 1.056 |
| gw -t4 | 200000 | 0.033 | 0.143 | 1.002 | 0.09 | 0.053 | 1.007 | 0.075 | 0.059 | 1.17 |
| gw -t4 | 2000000 | 0.054 | 0.173 | 0.887 | 0.09 | 0.084 | 0.791 | 0.16 | 0.059 | 2.148 |
| igv | 2000 | 0.029 | 3.613 | 36.748 | 3.377 | 0.236 | 26.977 | 0.341 | 0.252 | 5.701 |
| igv | 20000 | 0.03 | 4.224 | 40.718 | 3.377 | 0.847 | 59.758 | 0.469 | 0.252 | 7.801 |
| igv | 200000 | 0.038 | 8.43 | 59.207 | 3.377 | 5.054 | 95.666 | 0.805 | 0.252 | 12.575 |
| igv | 2000000 | 0.123 | 51.663 | 264.443 | 3.377 | 48.286 | 456.376 | 4.867 | 0.252 | 65.152 |
| igv -t4 | 2000 | 0.031 | 3.628 | 36.9 | 3.377 | 0.251 | 28.686 | 0.314 | 0.252 | 5.261 |
| igv -t4 | 20000 | 0.031 | 4.208 | 40.567 | 3.377 | 0.832 | 58.652 | 0.383 | 0.252 | 6.372 |
| igv -t4 | 200000 | 0.033 | 8.363 | 58.736 | 3.377 | 4.986 | 94.396 | 1.029 | 0.252 | 16.072 |
| igv -t4 | 2000000 | 0.054 | 50.524 | 258.613 | 3.377 | 47.147 | 445.611 | 5.663 | 0.252 | 75.799 |
| jb2export | 2000 | 0.029 | 2.829 | 28.773 | 2.668 | 0.161 | 18.423 | 0.493 | 0.445 | 8.252 |
| jb2export | 20000 | 0.03 | 3.066 | 29.559 | 2.668 | 0.399 | 28.129 | 0.53 | 0.445 | 8.82 |
| jb2export | 200000 | 0.038 | 5.192 | 36.463 | 2.668 | 2.524 | 47.785 | 0.947 | 0.445 | 14.799 |
| jb2export | 2000000 | 0.123 | 27.661 | 141.59 | 2.668 | 24.994 | 236.23 | 4.701 | 0.445 | 62.925 |
| samplot | 2000 | 0.029 | 0.678 | 6.891 | 0.57 | 0.108 | 12.281 | 0.127 | 0.117 | 2.117 |
| samplot | 20000 | 0.03 | 0.748 | 7.208 | 0.57 | 0.178 | 12.538 | 0.131 | 0.117 | 2.177 |
| samplot | 200000 | 0.038 | 1.304 | 9.16 | 0.57 | 0.734 | 13.902 | 0.158 | 0.117 | 2.472 |
| samplot | 2000000 | 0.123 | 7.425 | 38.005 | 0.57 | 6.855 | 64.789 | 0.536 | 0.117 | 7.169 |
| genomeview | 2000 | 0.029 | 0.634 | 6.452 | 0.231 | 0.404 | 46.078 | 0.143 | 0.14 | 2.392 |
| genomeview | 20000 | 0.03 | 3.926 | 37.849 | 0.231 | 3.696 | 260.665 | 0.144 | 0.14 | 2.403 |
| genomeview | 200000 | 0.038 | 34.407 | 241.645 | 0.231 | 34.176 | 646.969 | 0.157 | 0.14 | 2.453 |
| genomeview | 2000000 | 0.123 | 401.238 | 2053.8 | 0.231 | 401.007 | 3790.13 | 0.258 | 0.14 | 3.449 |

### Slide 7
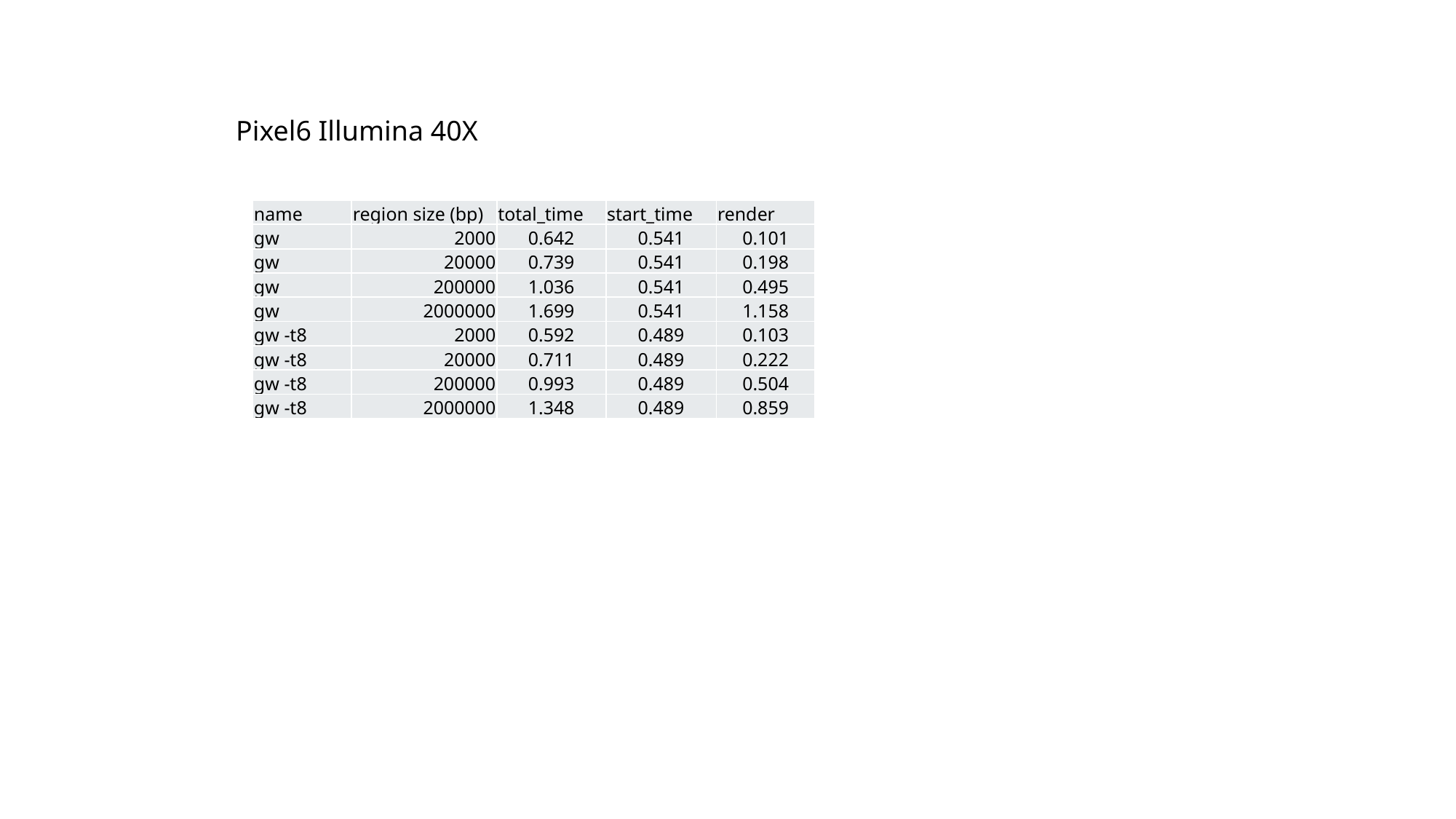

Pixel6 Illumina 40X
| name | region size (bp) | total\_time | start\_time | render |
| --- | --- | --- | --- | --- |
| gw | 2000 | 0.642 | 0.541 | 0.101 |
| gw | 20000 | 0.739 | 0.541 | 0.198 |
| gw | 200000 | 1.036 | 0.541 | 0.495 |
| gw | 2000000 | 1.699 | 0.541 | 1.158 |
| gw -t8 | 2000 | 0.592 | 0.489 | 0.103 |
| gw -t8 | 20000 | 0.711 | 0.489 | 0.222 |
| gw -t8 | 200000 | 0.993 | 0.489 | 0.504 |
| gw -t8 | 2000000 | 1.348 | 0.489 | 0.859 |
